# MBNL1 regulates programmed postnatal switching between regenerative and differentiated cardiac states

**DOI:** 10.1101/2023.03.16.532974

**Authors:** Logan R.J. Bailey, Darrian Bugg, Isabella M. Reichardt, C. Dessirée Ortaç, Jagadambika Gunaje, Richard Johnson, Michael J. MacCoss, Tomoya Sakamoto, Daniel P. Kelly, Michael Regnier, Jennifer M. Davis

## Abstract

Discovering determinants of cardiomyocyte maturity and the maintenance of differentiated states is critical to both understanding development and potentially reawakening endogenous regenerative programs in adult mammalian hearts as a therapeutic strategy. Here, the RNA binding protein Muscleblind-like 1 (MBNL1) was identified as a critical regulator of cardiomyocyte differentiated states and their regenerative potential through transcriptome-wide control of RNA stability. Targeted MBNL1 overexpression early in development prematurely transitioned cardiomyocytes to hypertrophic growth, hypoplasia, and dysfunction, whereas loss of MBNL1 function increased cardiomyocyte cell cycle entry and proliferation through altered cell cycle inhibitor transcript stability. Moreover, MBNL1-dependent stabilization of the estrogen-related receptor signaling axis was essential for maintaining cardiomyocyte maturity. In accordance with these data, modulating MBNL1 dose tuned the temporal window of cardiac regeneration, where enhanced MBNL1 activity arrested myocyte proliferation, and MBNL1 deletion promoted regenerative states with prolonged myocyte proliferation. Collectively these data suggest MBNL1 acts as a transcriptome-wide switch between regenerative and mature myocyte states postnatally and throughout adulthood.

## Introduction

The mammalian heart fundamentally changes during the transition between prenatal and postnatal development, which is termed cardiac maturation. During this process cardiomyocytes lose their fetal state and become terminally differentiated to support the circulatory demands on the adult heart. As cardiomyocytes terminally differentiate, they fundamentally change how energy is produced and used through the coordinated switching between embryonic and adult isoforms of metabolic, contractile, and ion handling genes^1^. In parallel, the primary mechanism of cardiomyocyte growth fundamentally switches from clonal expansion to hypertrophy^1^. Despite having considerable knowledge of the factors controlling early heart development, there is still a nascent understanding of the mechanisms that establish and maintain terminally differentiated cardiomyocyte states in the fully matured heart. Beyond furthering knowledge of cardiac development, delineating the mechanistic factors that control this transition are essential for rationally designing novel regenerative medicine approaches that address heart disease, as maintenance of cardiomyocyte terminal differentiation following injury underlies the negligible regenerative capacity of the mammalian heart following injury or stress^2–6^. Indeed, in model systems where cardiac regeneration occurs (e.g. zebrafish, neonatal mice), cardiomyocytes must first lose their terminally differentiated characteristics and return to a fetal transcriptomic state in order to proliferate^2–4,6–8^. It stands to reason that targeted destabilization of mature cardiomyocyte states could be used to promote endogenous heart regeneration.

Following terminal differentiation, what maintains a cardiomyocyte’s mature state is poorly understood but vital to the heart’s regenerative potential. Proliferative states and regenerative outcomes can be achieved in adult cardiomyocytes by modulating developmental signals like Hippo/YAP or ERBB2^9–14^ or reverting adult cardiomyocytes to a stem-like state with Yamanaka factors^15^. Yet, prolonged activation of these pathways can produce deleterious effects on cardiac function. Hence, we reasoned that controlling native factors that maintain mature myocyte states could improve the heart’s regenerative potential without reprogramming the cell. Surprisingly, few if any factors have been identified as essential for maintaining these terminally differentiated states. To date most studies have focused on inducers of cardiac maturation with an eye towards myogenic transcription factors^16–21^, epigenetic mechanisms^22–27^, and micro- and non-coding RNAs^28–30^. Despite the powerful influence of post-transcriptional regulation on cell behavior, little attention has been paid to mechanisms controlling RNA fate (alternative splicing, polyadenylation, stability, etc.), which ultimately shapes the myocyte’s transcriptional and proteomic landscape^31–33^. RNA binding proteins, like muscleblind-like 1 (MBNL1), exert global control over these regulatory processes yet they have not been widely interrogated in the context of cardiomyocyte terminal differentiation and regenerative programs. In the heart, MBNL1 expression promotes key fetal-to-adult splicing transitions for contractile, matrix, and calcium-handling proteins, and its expression rises and then plateaus in tandem with the timing of postnatal myocyte maturation^34–36^, suggesting that MBNL1 activity could be a key regulator of mature myocyte states. Indeed, MBNL1 promotes and maintains differentiated cell states in skeletal muscle, fibroblast, and blood lineages^36–41^, while its expression in stem cells minimizes their pluripotency^42,43^. These findings open the possibility that MBNL1 has similar functions in adult cardiomyocytes. Hence, we tested the hypothesis that MBNL1’s promotion and maintenance of mature states impairs cardiac regeneration.

Here, we demonstrate MBNL1 is essential for maintaining cardiomyocyte terminal differentiation through its transcriptome-wide stabilization of RNAs that drive maturation programs. Additionally, targeted MBNL1 overexpression early in development antagonized postnatal cardiomyocyte proliferation by stabilizing multiple cell cycle inhibitor transcripts and caused cardiac dysfunction due to early cardiomyocyte cell cycle exit and cardiac hypoplasia. Conversely, MBNL1 loss of function allowed for increased cardiomyocyte cell cycle entry and proliferation. Tuning the maturity of cardiomyocytes through MBNL1 modulation altered cardiac regenerative potential, whereby MBNL1 overexpression was sufficient to prevent cardiac regeneration in neonatal mice and MBNL1 loss of function extended the neonatal regeneration window.

## Results

### The kinetics of MBNL1 expression coincides with postnatal myocyte maturation

To independently identify drivers of cardiomyocyte terminal differentiation, neonatal (postnatal day 1, P1), juvenile (P7), and adult (P60) cardiac ventricles were subjected to RNA sequencing (RNAseq) (**Fig 1A**). Principal component analysis (PCA) of ventricle transcriptomes revealed time-dependent shifts in transcriptional space over the course of development (**Fig 1B**). To identify functional modules of genes which change over this period, gene-gene correlations were calculated for 13,290 genes which were differentially expressed in at least one comparison and K-nearest neighbor clustering was performed on the resulting gene-gene correlations (**Fig 1C-D**). The resulting 8 gene clusters were significantly enriched for functional categories known to be differentially regulated over the course of postnatal cardiac development. This analysis recapitulated known developmental changes including a postnatal decrease in proliferation and Yap/ErbB/Wnt signaling as well as an increase in mitochondrial metabolism and mature sarcomeric isoforms (**Fig 1E-L**). Additional modules identified included changes in proteostasis/control of translation, MAPK/PI3K-Akt signaling, and ECM/Focal Adhesions (**Fig S1A-H**). To identify regulators of cardiomyocyte terminal differentiation, clusters 3 and 4, which were consistently upregulated over the course of postnatal development, were more closely examined (**Fig 1I-L**). Both clusters 3 and 4 largely defined genes involved in mitochondrial structure/function, while cluster 4 also consisted of genes involved in mature mitochondrial and sarcomeric function which undergo fetal to adult isoform switching like Acsl1, Tnni3, and Myh6 (**Fig 1L**). This, coupled with the observation that cluster 4 expression begins to rise at P7 prior to the rise in cluster 3 expression, led us to focus on cluster 4 as a key gene regulatory network driving the establishment of cardiomyocyte terminal differentiation (**Fig 1K**). Known transcriptional and posttranscriptional regulators of postnatal cardiomyocyte maturation were grouped in cluster 4, including estrogen-related receptor (ERR) signaling (Esrra)^44,45^, retinoic acid (Rarb/Rarg) and retinoid-x (Rxrb/Rxrg) receptor signaling, as well as the developmentally regulated RNA binding proteins (RBPs) muscleblind-like protein (Mbnl1/Mbnl2) and Rbfox1 (**Fig 1M, S1I-O**). As we had previously identified MBNL1 as a central regulator of fibroblast differentiation and state maintenance^37,39^, MBNL1 was explored for its role in controlling cardiomyocyte terminal differentiation. RNAseq showed that MBNL1 gene expression consistently increases over the course of postnatal cardiac development as expression of its counter-regulatory RBP CELF1 decreases (**Fig 1M-N**). At the protein level, MBNL1 expression rapidly increases from P1 to P4 before peaking around P10 and gradually falling back to a lower maintenance level in P60 ventricles (**Fig 1O**). This increase in expression occurs during the same time frame as known and predicted changes occur in other facets of cardiomyocyte terminal differentiation. Notably, as MBNL1 expression increases there are time-dependent decreases in proliferation (AURKB), YAP (phosphorylated-YAP and total YAP), and glycolytic proteins (GAPDH), time-dependent increases in mitochondrial oxidative phosphorylation components (Complex I, II, III, V), and time-dependent isoform switches between fetal (ssTnI) and adult cardiac (cTnI) sarcomeric isoforms (**Fig 1O**). Collectively, this analysis suggests MBNL1 drives and maintains the differentiated state in cardiomyocytes.

**Figure 1.**
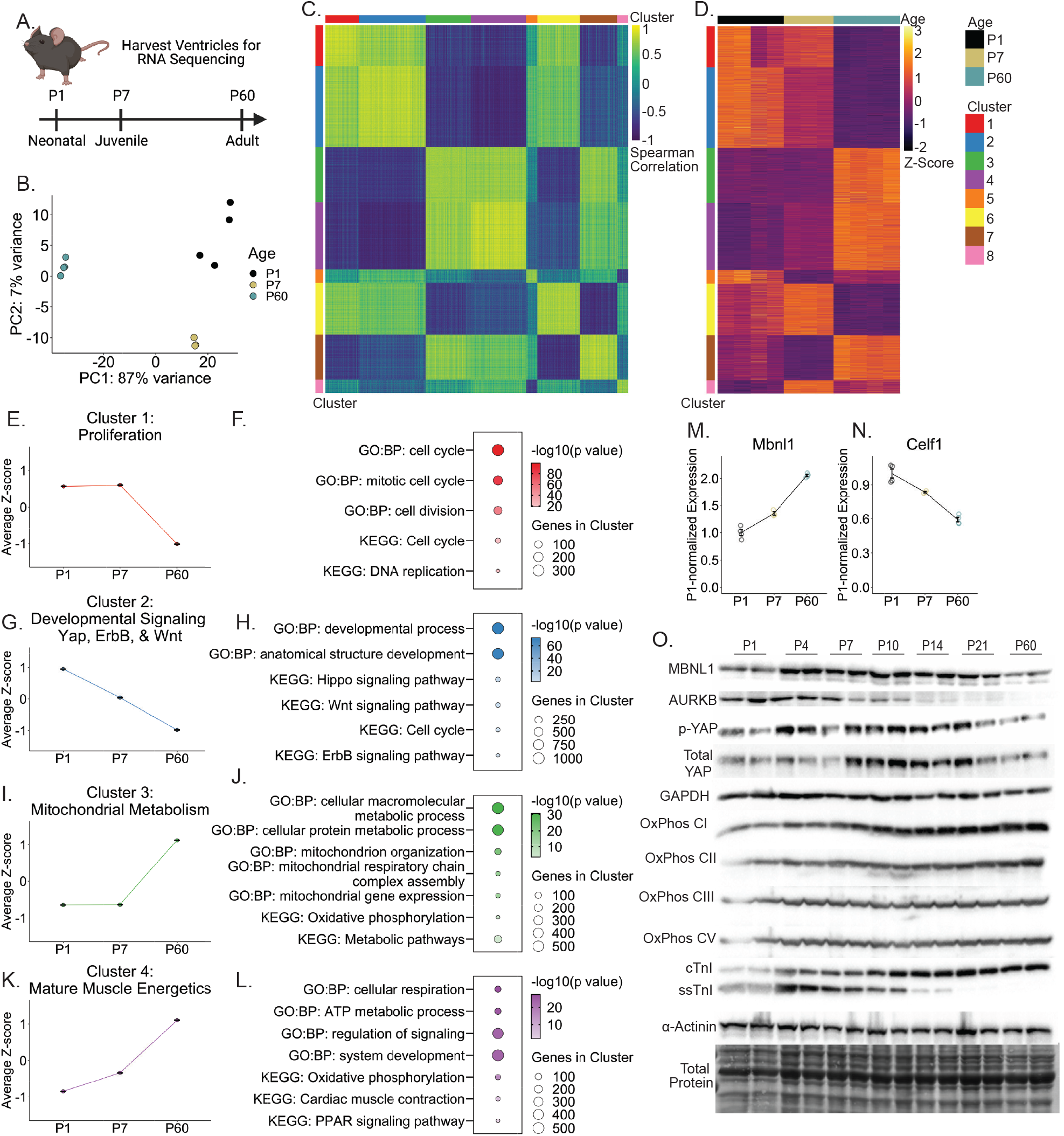
The kinetics of MBNL1 expression coincides with postnatal myocyte maturation. (A) Experimental outline for RNA sequencing of wild type ventricles at postnatal day 1 (P1, N=4), 7 (P7, N=3), and 60 (P60, N=4). (B) Principal component analysis of ventricle transcriptomes. (C) Pairwise gene-gene spearman correlations for all differentially expressed genes (DEG) color coded by k-nearest neighbor (KNN) clustering. (D) Normalized gene expression of all DEG color coded by age and KNN clustering. (E,G,I,K) Mean normalized expression ± SEM of genes within each cluster over time. (F,H,J,L) Representative Gene Ontology: Biological Processes (GO:BP) and Kyoto Encyclopedia of Genes and Genomes (KEGG) terms for each cluster. (M-N) P1-normalized expression of Mbnl1 (M) and Celf1 (N) by RNAseq over time. Dots represent biological replicates. Lines show mean ± SEM. (O) Western blots of heart lysates at postnatal days 1 (P1), 4, 7, 10, 14, 21, and 60 for MBNL1, Aurora B Kinase (AURKB), phosphorylated YAP (p-YAP), total YAP (YAP), GAPDH, oxidative phosphorylation complex I, II, III, V (OxPhos Complex I/II/III/V), cardiac troponin I (cTnI), slow skeletal troponin I (ssTnI), α-Actinin, and total protein. α**-**Actinin and total protein were used as loading controls. N=2 for each time point. See also Figure S1.

### MBNL1 is necessary to maintain the terminally differentiated state of adult cardiomyocytes

A mouse model with inducible loss of MBNL1 function solely in cardiomyocytes (MBNL1 iKO) was used to examine whether MBNL1 is necessary for maintaining cardiomyocyte terminal differentiation. Here, conditional MBNL1 knockout (MBNL1^F/F^)^37^ mice were crossed with mice containing a tamoxifen-inducible Cre driver expressed by the Myh6 promoter (Myh6^iCre^)^46^ (**Fig 2A**). Western blotting of phenylephrine-treated neonatal ventricular myocytes (NVMs) showed robust deletion of MBNL1 in cardiomyocytes following adenoviral Cre induction relative to βgal control (**Fig 2B**). Within 14 days of tamoxifen induction, cardiomyocyte-specific MBNL1 deletion caused systolic dysfunction and left ventricular dilation relative to control animals (**Fig 2C-D**). To confirm that this phenotype was not due to tamoxifen toxicity echocardiography was repeated in the absence of tamoxifen chow (**Fig. S2A**). Additionally, a rescue experiment was performed by crossing a Cre-inducible MBNL1 Transgene (MBNL1 Tg)^39^ into the MBNL1 iKO mice allowing for simultaneous MBNL1 excision and overexpression (MBNL1 iKO+iOE) (**Fig. S2B**). Following tamoxifen induction, MBNL1 iKO mice had systolic dysfunction relative to Myh6^iCre^ mice, and MBNL1 overexpression was sufficient to rescue this phenotype (**Fig. S2A**). MBNL1 iKO mice also had elevated heart weight/body weight ratios (**Fig 2E**) and elevated lung weight/body ratios relative to control littermates, indicating that loss of MBNL1 function rapidly transitions the heart to failure (**Fig 2F**).

**Figure 2.**
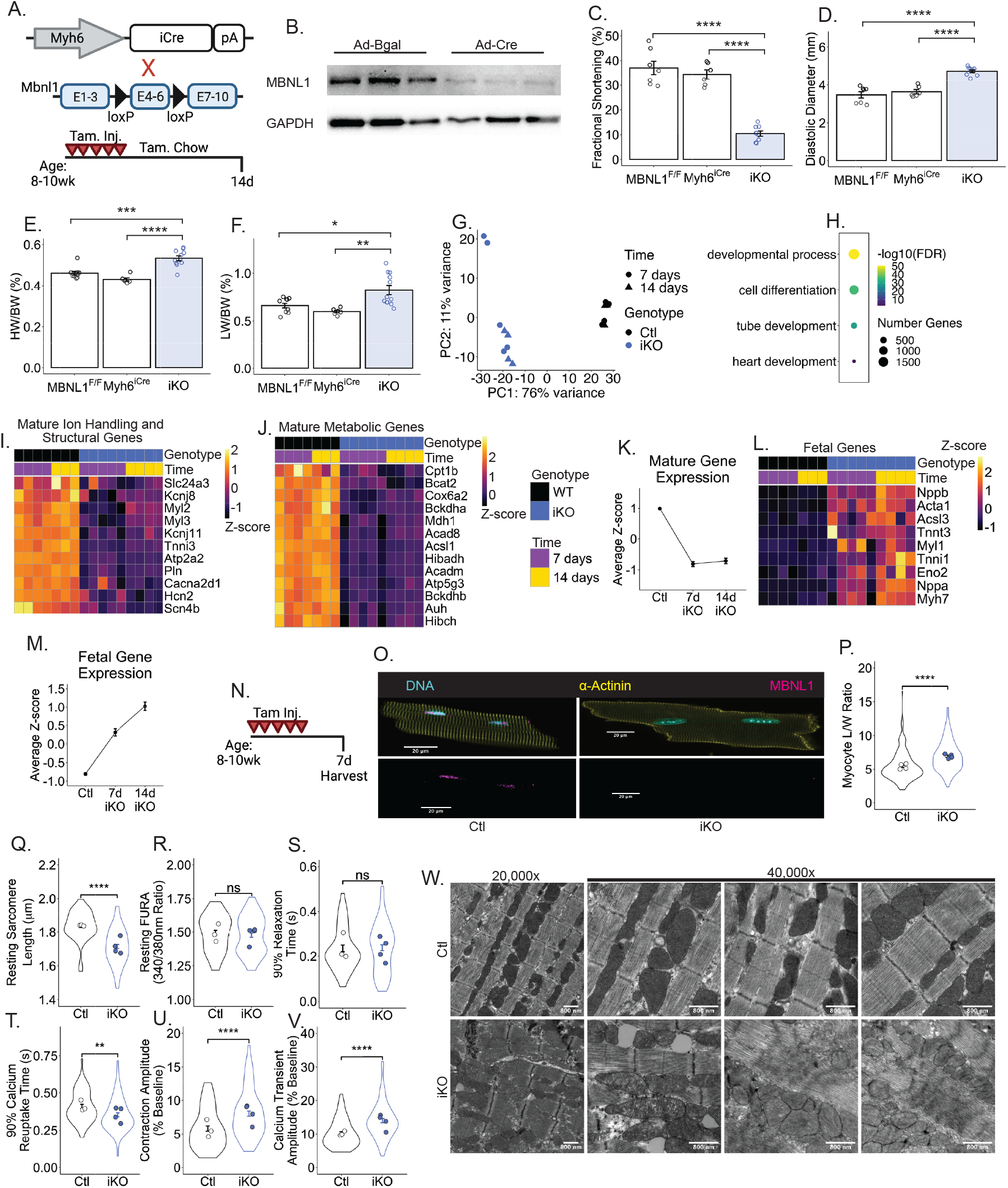
MBNL1 is necessary to maintain the terminally differentiated state of adult cardiomyocytes. (A) Breeding scheme for generating MBNL1 iKO mice (top) and experimental schematic (bottom). (B) Western blot of MBNL1 from MBNL1^F/F^ neonatal ventricular cardiomyocytes adenovirally transduced with βgal (N=3) or Cre (N=3) and treated with phenylephrine. GAPDH was used as a loading control. (C-D) Quantification of left ventricular fractional shortening (C) and diastolic diameter (D) in control and MBNL1 iKO mice by echocardiography (N=6-9). Dots are biological replicates. Bars are mean ± SEM. ****p<0.0001 (E) Heart weight/body weight ratio (N=6-12). Dots are biological replicates. Bars are mean ± SEM. ***p<0.001, ****p<0.0001. (F) Lung weight/body weight ratio (N=6-12). Dots are biological replicates. Bars are mean ± SEM. *p<0.05, **p<0.01. (G) Principal component analysis of ventricle transcriptomes from control and MBNL1 iKO mice at 7- (N=4-5) and 14-days (N=3-4) following tamoxifen initiation. (H) Dotplot of select GO:BP terms from gene ontology analysis of differentially expressed genes (DEG) by RNAseq. Color of dot represents Benjamini-Hochberg-corrected p-value (FDR). Size of dot is proportional to the number of differentially expressed genes in that category. (I-J) Heatmap of DEG representing mature cardiac ion-handling and structural transcripts (I) and mature cardiac metabolic transcripts (J). (K) Mean normalized expression ± SEM of mature transcripts by RNAseq. (L) Heatmap of DEG representing fetal cardiac transcripts. (M) Mean normalized expression ± SEM of fetal transcripts by RNAseq. (N) Tamoxifen dosing strategy for remainder of experiments (Figure 2O-W). (O) Representative immunocytochemical images of isolated cardiomyocytes stained with α-Actinin (Yellow) and MBNL1 (magenta) antibodies. (P) Myocyte length/width ratio quantification (n=200 myocytes/N=4 mice). Dots represent the mean for an individual mouse. Violin plots represent the distribution and error bars show the SEM of individual myocytes. Individual myocytes were compared between groups. ****p<0.0001. (Q-V) Quantification of isolated cardiomyocyte resting sarcomere length (Q), resting calcium by FURA ratio (R), time to 90% relaxation (S), time to 90% calcium reuptake (T), contraction amplitude (U), and calcium transient amplitude (V). Dots represent the mean for an individual mouse. Violin plots represent the distribution and error bars show the SEM of individual myocytes. **p<0.01, ****p<0.0001, ns=not significant. Control n=59 myocytes/N=3 mice, iKO n=84/N=4. (W) Representative electron micrographs from MBNL1 iKO (n=2) and control (n=2) hearts at 20,000x and 40,000x magnification. See also Figure S2.

To examine whether the deterioration in cardiac structure-function was due to a MBNL1-dependent loss of mature myocyte transcriptome, RNAseq was performed on MBNL1 iKO and control (Ctl) ventricles. PCA analysis separately clustered MBNL1 iKO ventricle samples along the first principal component relative to controls regardless of time following excision (**Fig 2G**). MBNL1 iKO mice had broad dysregulation of cardiac developmental signaling (**Fig 2H**). Expression of mature metabolic and functional cardiac genes which canonically define cardiomyocyte maturity were uniformly downregulated in MBNL1 iKO mice, whereas fetal genes were upregulated relative to control (**Fig 2I-M**). With respect to the kinetics of these changes, mature gene expression was lost within 7 days of MBNL1 deletion and maintained over time (**Fig 2K**). Fetal gene expression, which could also represent nonspecific myocardial stress, gradually increased with time in MBNL1 knockouts (**Fig 2M**). Taken together, these kinetics suggest MBNL1 iKO mice first lose expression of mature gene isoforms that define their terminally differentiated state and then begin upregulating fetal isoform expression.

Early morphologic and ultrastructural changes were also observed in adult cardiomyocytes after MBNL1 loss of function (**Fig 2N**). Consistent with the organ-level phenotype, individual MBNL1 iKO myocytes underwent dilated remodeling relative to control cells (**Fig 2O-P, S2C-E**). Furthermore, functional analysis of excitation contraction (EC)-coupling suggested that MBNL1 iKO cardiomyocytes have increased calcium sensitivity, given diastolic sarcomere lengths were shorter despite similar resting calcium concentrations and relaxation kinetics were unchanged despite faster calcium reuptake (**Fig 2Q-T**). MBNL1 iKO myocytes also had increased shortening and calcium transient amplitudes, consistent with the dilated remodeling of the myocytes and increased calcium sensitivity (**Fig 2U-V**). Heightened calcium sensitivity is a hallmark of immature cardiomyocytes which could be explained by decreased cardiac troponin T (cTnT) exon 5 exclusion (a canonical MBNL1-regulated splice event ^47^) or an increased slow skeletal troponin I (ssTnI/Tnni1) to cardiac troponin I ratio (cTnI/Tnni3), both of which were observed in MBNL1 iKO mice relative to control via RNAseq (**Fig S2F-I**). Furthermore, consistent with the observed gene expression changes, MBNL1 iKO mice show ultrastructural defects in both mitochondria and sarcomeres by electron microscopy in which MBNL1 iKO hearts had disrupted mitochondrial cristae density throughout the tissue and focal patches of sarcomere disarray/disassembly (**Fig 2W**). Overall, this data demonstrates that loss of MBNL1 causes structural remodeling, functional changes, and loss of a mature transcriptome, all of which are consistent with a loss of cardiomyocyte terminal differentiation.

### MBNL1 regulates transcriptome-wide cardiomyocyte RNA stability

To determine how MBNL1 regulates cardiomyocyte terminal differentiation, MBNL1’s protein interactome was unbiasedly assessed by a proximity labeling assay using the promiscuous biotin ligase BioID2 (**Fig 3A**)^48^. Here, BioID2 was cloned onto the C-terminus of MBNL1. Correct localization of the fusion construct and biotin conjugation was verified by immunocytochemistry (ICC) (**Fig S3A**). Both the MBNL1-BioID2 conjugate and unconjugated BioID2 were adenovirally transduced into rat neonatal ventricular myocytes (NVMs) and the interactome labelled with biotin. Isolated biotinylated proteins were analyzed by liquid chromatography-mass spectrometry, which identified 229 significantly enriched MBNL1 protein interactors with virtually no overlap detected in the unconjugated BioID2 control (**Fig S3B-D**). Gene ontology analysis showed that MBNL1’s interactome is enriched for proteins involved in all facets of RNA metabolism, including mRNA splicing, mRNA degradation/stability, and regulation of translation (**Fig S3E**), with the RNA stability category containing 14% of the total interactome (32 MBNL1 interactors, **Fig 3B, S3F**). Several of these interactors are known to promote mRNA stability principally by binding transcripts’ 3’UTR (**Fig 3C**). Indeed, previous cross-linking and immunoprecipitation sequencing (CLIPseq) studies demonstrated that approximately half of MBNL1 binding sites are located in the 3’UTR of its target transcripts^49,50^. To further examine RNA stability as a potential mechanism, all the transcripts bound to MBNL1 in cardiomyocytes were identified using a native RNA immunoprecipitation followed by sequencing (RIPseq). Here, rat NVMs were adenovirally transduced with FLAG-tagged MBNL1 (**Fig 3D**) and immunoprecipitation with a FLAG antibody yielded a total of 4413 transcripts that were significantly enriched relative to the input control (**Fig 3E**). Significant overlap was found between transcripts identified in this RIPseq assay and those identified previously by similar approaches performed in other cell types^39,50^ (**Fig 3E, S4A**). Gene ontology analysis of the MBNL1-bound transcripts showed an enrichment for functions like cell differentiation and heart development, while non-MBNL1-bound transcripts were enriched for mitochondrially-derived, spliceosomal, and ribosomal transcripts (**Fig 3F, S4B**). Notably, transcripts for key cardiomyocyte specification (Gata4, Hand2, Tbx5, Tbx3, Nkx2-5) and maturation (Esrra, Rara, Rxra, Rxrb, and Srf) factors were bound by MBNL1 (**Fig 3G**). Additionally, nearly half of the transcripts bound by MBNL1 were differentially expressed in MBNL1 iKO ventricles (**Fig 3H**), which was surprising because MBNL1 is traditionally viewed as a regulator of alternative splicing in the heart^34–36,40,47^. Yet, we observed an order of magnitude more differentially expressed versus alternatively spliced genes in response to MBNL1 deletion (**Fig 3H**), which suggests MBNL1’s stability function is a primary determinant of its role in myocyte fate decision making.

**Figure 3.**
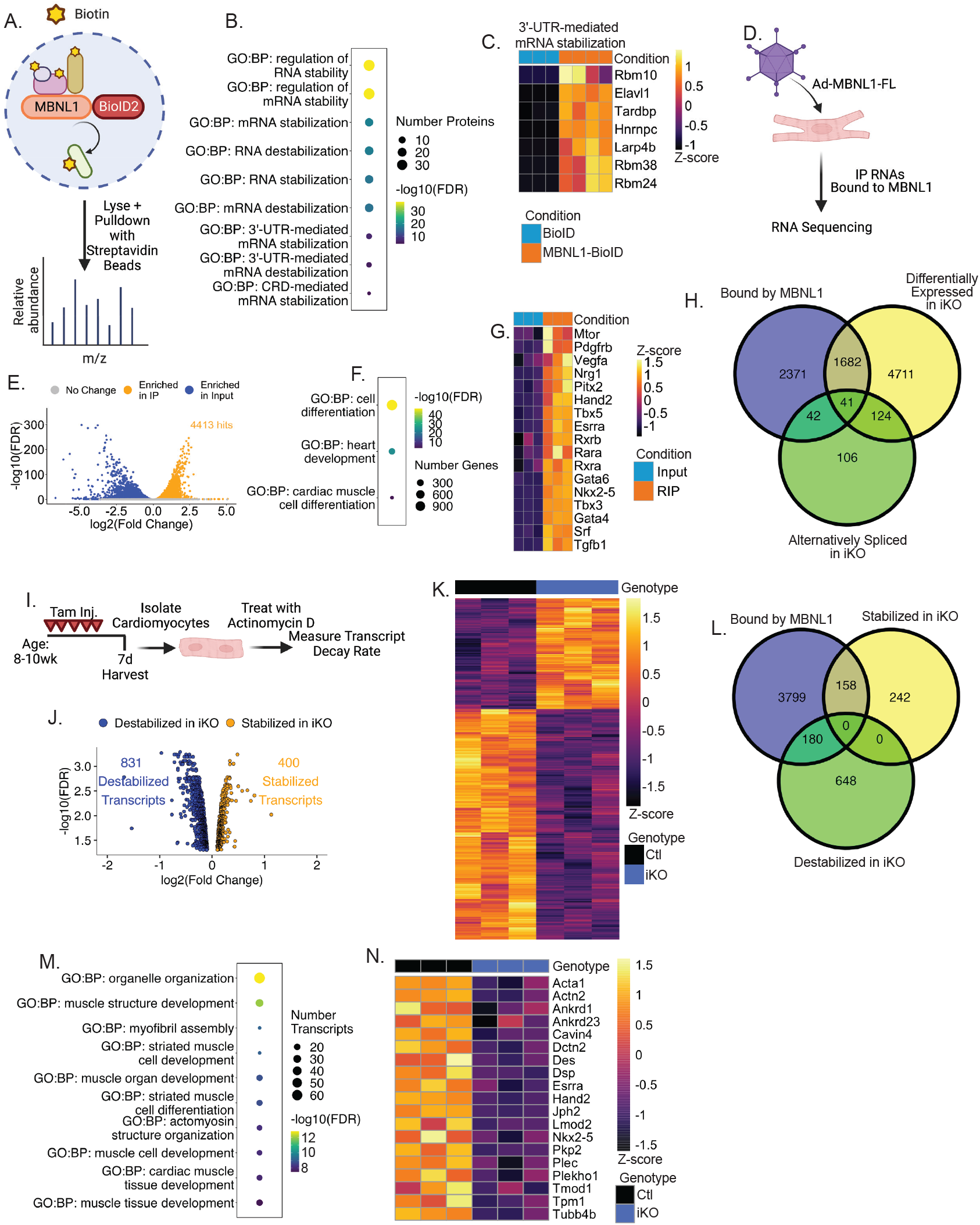
MBNL1 regulates cardiomyocyte transcriptome-wide RNA stability. (A) Experimental schematic of BioID proximity labeling experiment. (B) GO:BP terms relating to RNA stability from gene ontology analysis of MBNL1 protein interactors. (C) Heatmap of enrichment of MBNL1 protein interactors in the GO:BP category “3’-UTR-mediated mRNA stabilization” (N=3/4). (D) RIPseq experimental schematic. (E) Volcano plot of RIPseq results. (F) Select GO:BP terms from gene ontology analysis of transcripts enriched in MBNL1 RIP. (G) Heatmap of enrichment of select cardiomyogenic transcripts in input and RIP (N=3/3). (H) Venn diagram of genes bound by MBNL1, DEG in MBNL1 iKO ventricles, and alternatively spliced genes in MBNL1 iKO ventricles. (I) Experimental setup for Actinomycin D (ActD) pulse-chase transcript stability assay. (J) Volcano plot of ActD-sequencing experiment differential stability analysis. (K) Heatmap showing normalized fraction of RNA remaining 8 hours following ActD treatment for all differentially stabilized transcripts between control and MBNL1 iKO mice (N=3/3). (L) Venn diagram comparing transcripts bound by MBNL1, transcripts stabilized in MBNL1 iKO, and transcripts destabilized in MBNL1 iKO. (M) Top 10 GO:BP terms of transcripts bound by MBNL1 and destabilized in MBNL1 iKO. (N) Heatmap showing normalized fraction of RNA remaining 8 hours following ActD treatment for representative transcripts bound by MBNL1 and destabilized in MBNL1 iKO (N=3/3). See also Figure S3 and Figure S4.

To determine if MBNL1-dependent regulation of RNA stability underlies the maturation program in cardiomyocytes, isolated MBNL1 iKO and control myocytes were treated with the transcription inhibitor Actinomycin D (ActD, 2μM), and transcriptome-wide transcript decay rates were measured by RNAseq 4 and 8 hours after treatment (**Fig 3I**). In MBNL1 iKO myocytes 1,231 transcripts had differential transcript stability with more than 65% of them destabilized (**Fig 3J-K**). Of the destabilized transcripts, 180 were bound by MBNL1 (**Fig 3L**). Within the subset of transcripts that are both bound by MBNL1 and destabilized in MBNL1 iKO were factors vital to cardiomyocyte maturity and function, including structural transcripts like Acta2, Dsp, Jph2, and Tpm1 as well as cardiomyogenic transcription factors like Nkx2.5, Hand2, and Esrra (**Fig 3M-N**). Taken together, these data suggest that MBNL1 promotes and maintains mature cardiomyocyte states at least in part through its RNA stabilizing function.

### MBNL1-dependent stabilization of estrogen-related receptor signaling maintains the terminally differentiated state of adult cardiomyocytes

Given that MBNL1 binds many important cardiomyocyte fate specification and maturation transcripts and that it controls transcriptome-wide RNA stability, we reasoned that MBNL1 regulates cardiomyocyte terminal differentiation through stabilization of specific cardiomyocyte maturation transcripts. To determine which of the 4,413 transcripts bound by MBNL1 in cardiomyocytes are critical to maintaining their mature state, our data sets were searched for genes that satisfied the following criteria: (1) genes downregulated in MBNL1 iKO hearts, (2) transcription factors predicted to explain downregulated genes, (3) transcripts bound by MBNL1, and (4) transcripts with MBNL1-dependent stability. This search uncovered one intersecting gene: the estrogen-related receptor (ERR) transcript Esrra (**Fig 4A**). The ERRs are critical signaling mediators of postnatal cardiac maturation which simultaneously promote mitochondrial and sarcomeric maturation and, importantly, Esrra and Mbnl1 gene expression rise over a similar developmental timeframe^44,45^. Additionally, Esrra was clustered with MBNL1 in our terminal differentiation driver cluster and its upregulation slightly lags increased MBNL1 expression in maturing cardiomyocytes (**Fig 1M, S1I**). Critically, Esrra and both of its paralogs Esrrb and Esrrg showed decreased expression in MBNL1 iKO mice compared to control (**Fig 4B**). Additionally, direct MBNL1 binding of Esrra and Esrrg (the two major cardiac paralogs) was confirmed by RIP-qPCR (**Fig 4C**). qPCR confirmation of the ActD transcript stability assay revealed that MBNL1 is necessary for Esrra transcript stabilization but not for Esrrg (**Fig 4D-E**). Consistent with the destabilization and downregulation of the Esrra transcript, ERR signaling is broadly inhibited in MBNL1 iKO ventricles relative to control, which is demonstrated by the global downregulation of ERR-activated genes and reciprocal activation of ERR-suppressed genes (**Fig 4F-G**). However, ERR signaling can also be inhibited due to poor heart function^51^. To further confirm that MBNL1 directly regulates ERR signaling through stabilizing the Esrra transcript in this context versus a secondary change from cardiac failure, isolated wild type adult rat cardiomyocytes were adenovirally transduced with a short hairpin RNA (shRNA) targeting MBNL1 (shMBNL1) or non-targeting shRNA control (shLacZ) for 5 days. MBNL1 knockdown produced robust downregulation of MBNL1, as well as a shift towards fetal isoforms like a decreased Tnni3/Tnni1 ratio as observed in our MBNL1 iKO model (**Fig 4H-I**). Additionally, MBNL1 knockdown produced downregulation of the Esrra transcript as well as downregulation of key downstream ERR signaling targets like Acsl1 and Esrrg, validating that MBNL1 regulates this signaling pathway (**Fig 4J-L**). Furthermore, simultaneous MBNL1 knockdown and human ERR-alpha (ERRα) overexpression rescues these transcriptional changes (**Fig 4J-L**). Notably, overexpression of human ERRα, which should drive increased rat Esrra expression, produced less Esrra in shMBNL1 treated cells compared to control, further emphasizing the important regulatory role of MBNL1 in stabilizing the Esrra transcript (**Fig 4J**). Together, this data suggests that MBNL1 creates a gain in Esrra function and its canonical downstream signaling pathway to maintain cardiomyocyte terminal differentiation.

**Figure 4.**
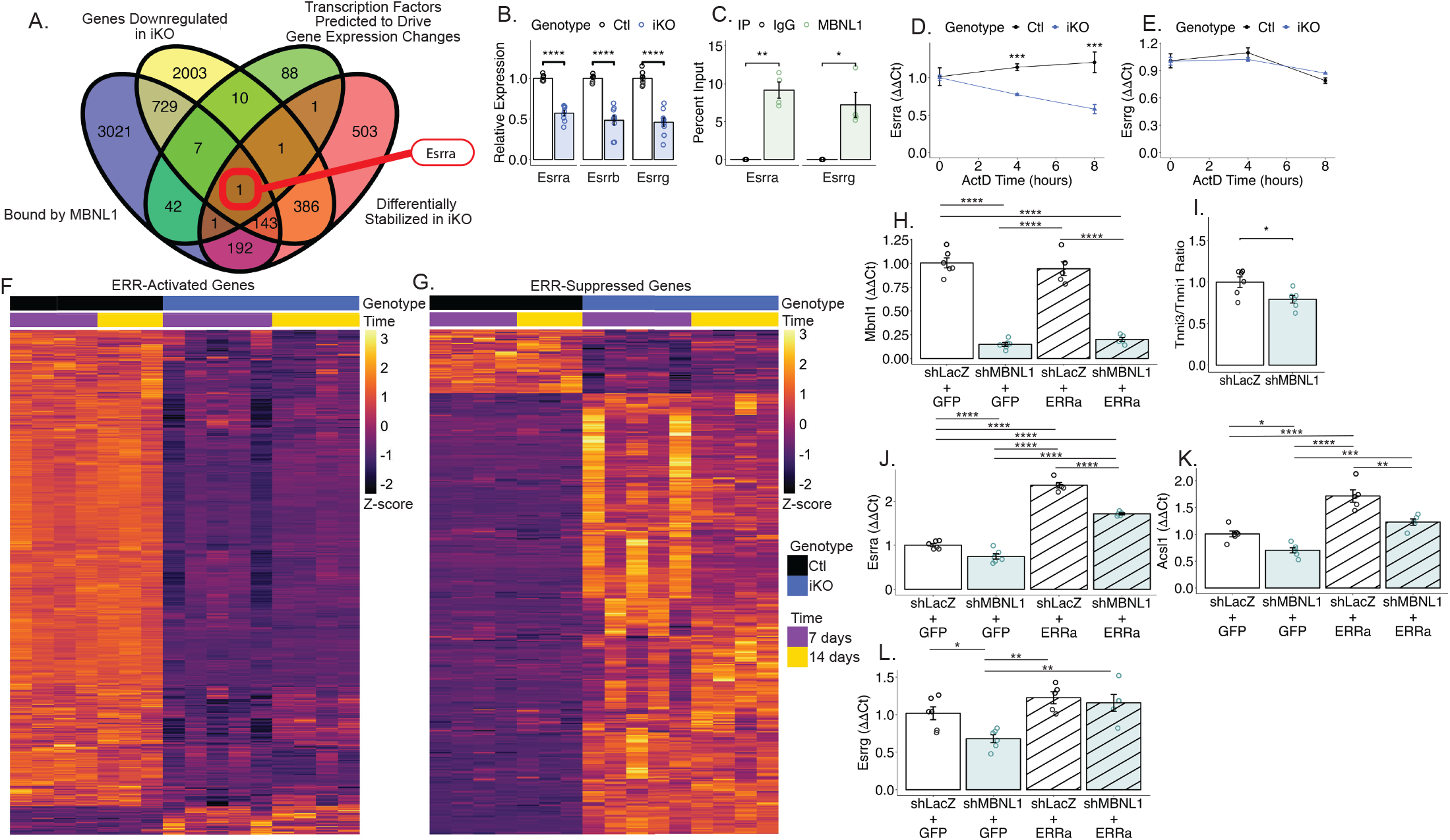
MBNL1-dependent stabilization of Estrogen-related Receptor signaling maintains the terminally differentiated state of adult cardiomyocytes. (A) Venn diagram comparing transcripts downregulated in MBNL1 iKO mice, transcription factors predicted to drive expression of these downregulated genes, transcripts bound by MBNL1, and differentially stabilized transcripts in MBNL1 iKO mice. (B) Expression of Esrra, Esrrb, and Esrrg by RNAseq (N=7/9). Dots represent biological replicates. Bars represent mean ± SEM. ****p<0.0001. (C) RIP-qPCR for Esrra and Esrrg in IgG versus MBNL1 immunoprecipitation (N=4/4). *p<0.05, **p<0.01. (D-E) qPCR quantification of Esrra (D) and Esrrg (E) expression following actinomycin D (ActD) treatment (N=3/3). Dots represent mean ± SEM. ***p<0.001. (F-G) Heatmap of ERR-activated gene expression (F) and ERR-suppressed gene expression (G) in control and MBNL1 iKO ventricles by RNAseq (N=7/9). (H-L) Expression of Mbnl1 (H), Tnni3/Tnni1 ratio (I), Esrra (J), Acsl1 (K), and Esrrg (L) by qPCR in wild type adult rat ventricular cardiomyocytes transduced with non-targeting shRNA (shLacZ), MBNL1-targeting shRNA (shMBNL1), GFP, and/or human ERR-alpha (ERRa) adenoviruses (N=5-6). Dots represent biological replicates. Bars represent mean ± SEM. *p<0.05, **p<0.01, ***p<0.001, ****p<0.0001.

### Early MBNL1 overexpression antagonizes postnatal cardiomyocyte proliferation

As MBNL1 was necessary to maintain cardiomyocyte terminal differentiation, we next sought to determine if overexpression of MBNL1 could potentiate maturation of postnatal hearts, resulting in either early switching between hyperplastic and hypertrophic growth or exaggerated maturation and hypertrophic growth. To test this hypothesis, a cardiomyocyte-specific MBNL1 transgenic mouse (MBNL1 cOE) was generated by mating a conditional MBNL1 transgene (MBNL1 Tg)^39^ with a constitutively active cardiomyocyte Cre driver (Myh6^Cre^)^52^ (**Fig 5A**). Western blotting of P1 ventricles showed strong overexpression of MBNL1 relative to littermate control mice which was maintained into adulthood (**Fig 5B-C**). Rather than accelerating maturation and promoting hypertrophic cardiac growth, young adult (P60) MBNL1 cOE mice had both systolic and diastolic dysfunction (**Fig 5D-F**) that was in part due to reduced cardiomyocyte shortening and calcium transient amplitudes as well as slower kinetics (**Fig S5A-H**). In addition, MBNL1 cOE mice had significantly smaller ventricle weight/body weight ratios despite having larger cardiomyocyte cross-sectional areas which is indicative of hypertrophic growth (**Fig 5G-J**). When compared to control littermates, cardiomyocytes isolated from MBNL1 cOE mice had increased areas and length/width (L/W) ratios driven by increased width, a morphological hallmark of parallel myofiber addition and concentric hypertrophy (**Fig 5K, S5I-K**). These disparate phenotypes of smaller ventricles with hypertrophic cardiomyocytes suggested there were fewer total cells in MBNL1 cOE hearts. Consistent with this hypothesis, fewer total cardiomyocytes were isolated from MBNL1 cOE hearts relative to control littermates despite negligible rates of cardiomyocyte death during the isolation procedure (**Fig 5L, Fig S5L**). Given MBNL1 negatively regulates proliferation of other cell types and acts as a tumor suppressor in cancer^53–55^, we hypothesized that MBNL1 overexpression suppresses postnatal cardiomyocyte proliferation. To test this hypothesis, myocardial sections were analyzed by immunohistochemistry at postnatal day 8 during the last wave of postnatal cardiomyocyte proliferation^56^. Here, both cardiomyocytes and non-cardiomyocytes were genetically labeled using a Cre-dependent dual fluorescent reporter (mTmG)^57^. All cells in mTmG mice express a membrane-targeted tdTomato (mT) unless Cre activity excises the mT and moves a membrane-targeted eGFP (mG) sequence in frame for expression. This permits simultaneous labeling of Myh6 lineage cardiomyocytes (mG+) and non-cardiomyocytes (mT+). Analysis of lineage-traced cardiomyocytes confirmed that MBNL1 cOE hearts had decreased cardiomyocyte-specific cell cycle activity relative to control as demonstrated by decreased EdU incorporation, Ki67, and phospho-histone H3 (pH3) staining (**Fig 5M-S**). Moreover, RNAseq analysis of P7 ventricles demonstrated that genes differentially expressed between MBNL1 cOE and controls were highly functionally enriched for cell cycle regulation based on gene ontology analysis (**Fig 5T**). Consistent with the cell cycle imaging, MBNL1 cOE ventricles had uniform downregulation of positive cell cycle regulators like Ccne1, Ccnb1, and Aurkb, as well as upregulation of negative cell cycle regulators like Cdkn1a and Gadd45g (**Fig 5U**). To ensure that this reduction in cardiomyocyte proliferation contributes to the cardiac dysfunction observed in MBNL1 cOE mice, MBNL1 Tg mice were crossed with Myh6^iCre^ mice (MBNL1 iOE) to control the timing of MBNL1 overexpression (**Fig S6A**). Here, MBNL1 overexpression was initiated at 8-10 weeks of age, after cardiomyocytes have largely exited the cell cycle (**Fig S6B**), and cardiac structure and function were examined over the course of 30 days. Ventricle weight/body weight ratios did not differ between genotypes (**Fig S6C-D**), and MBNL1 iOE caused mild systolic impairment following tamoxifen treatment with no diastolic dysfunction or structural remodeling by 30 days (**Fig S6E-M**). Together, this indicates that (1) the dysfunction seen in MBNL1 cOE mice is due to inhibition of postnatal cardiomyocyte proliferation rather than acute effects of MBNL1 overexpression in fully mature cardiomyocytes and (2) early MBNL1 overexpression prematurely halts postnatal cardiomyocyte proliferation, leading to cardiomyocyte hypoplasia.

**Figure 5.**
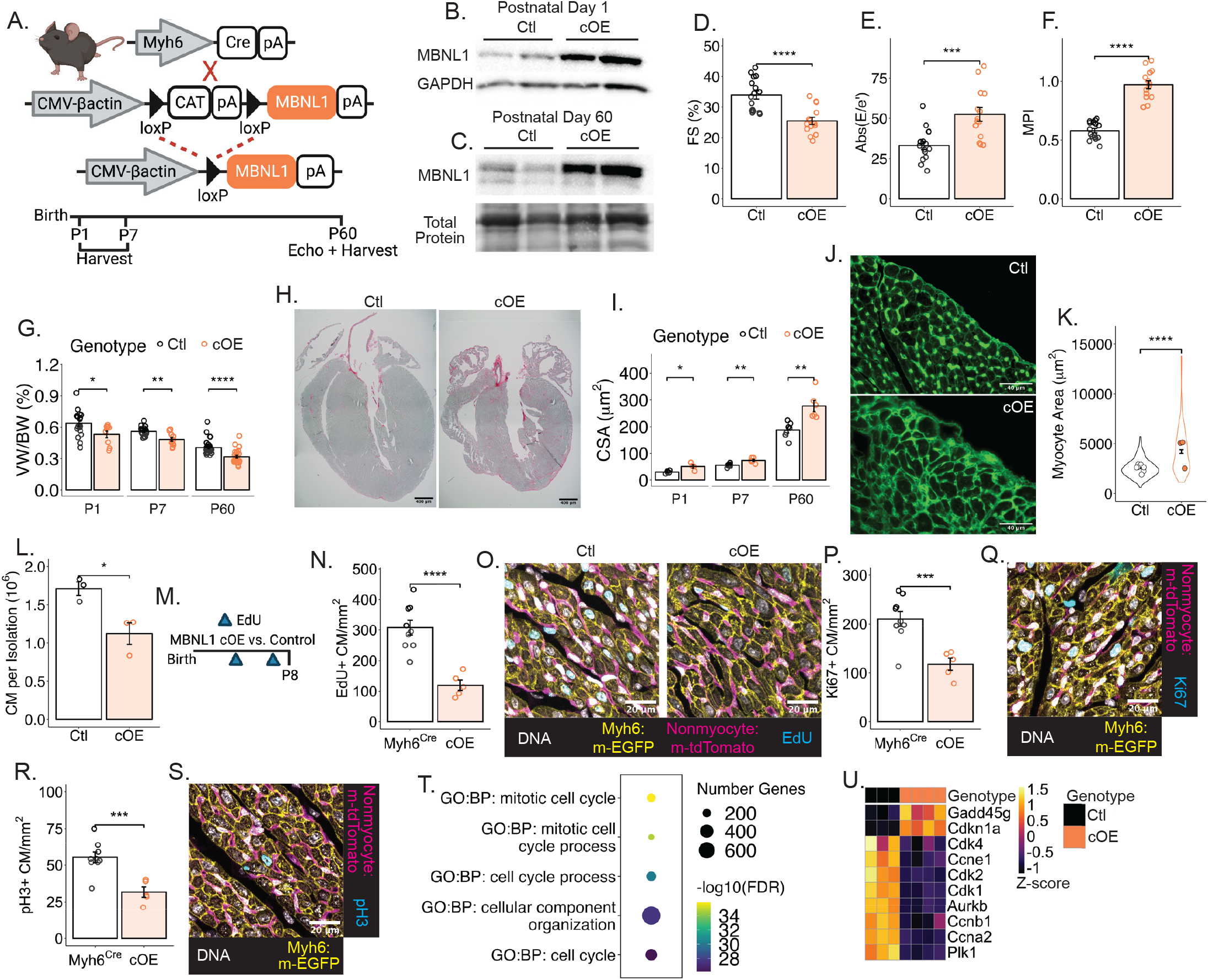
Early MBNL1 overexpression antagonizes postnatal cardiomyocyte proliferation. (A) Breeding schematic to generate MBNL1 cOE mice (top) and experimental timeline (bottom). (B-C) Western blotting for MBNL1 in P1 ventricles (B) and isolated P60 cardiomyocytes (C). GAPDH and total protein were used as loading controls (N=2/2). (D-F) Quantification of fractional shortening (FS) (D), absolute value of E/e’ ratio (Abs(E/e’) (E), and myocardial performance index (MPI) (F) by echocardiography (N=16/14). Dots represent biological replicates. Bars represent mean ± SEM. ***p<0.001, ****p<0.0001. (G) Ventricle weight/body weight (VW/BW) ratio of P1, P7, and P60 MBNL1 cOE mice versus controls (N=9-37). Dots represent biological replicates. Bars represent mean ± SEM. *p<0.05, **p<0.01, ****p<0.0001. (H) Representative images of sirius red/fast green stained myocardial sections showing smaller ventricles in P60 MBNL1 cOE mice compared to control. (I) Quantification of average cardiomyocyte cross sectional area (CSA) by wheat germ agglutinin (WGA) staining at P1, P7, and P60 (N=4-8). Dots represent biological replicates. Bars represent mean ± SEM. Mouse average CSA was compared between groups. *p<0.05, **p<0.01. (J) Representative image of WGA-stained myocardial sections at P60 for control and cOE hearts. (K) Quantification of area of isolated cardiomyocytes at P60. Dots represent the mean for an individual mouse. Violin plots represent the distribution and error bars show the SEM of individual myocytes. Individual myocytes were compared between groups. Control n=406 myocytes/N=8 mice, cOE n=146/N=3. ****p<0.0001 (L) Cardiomyocyte (CM) cell yield following cell isolation (N=3/3). *p<0.05. (M) EdU dosing schematic for neonatal mice. EdU is delivered at P4 and P7. (N-S) Quantification and representative images of EdU (N,O), Ki67 (P,Q), and pH3 (R,S) positive cardiomyocytes (CM) by immunohistochemistry at P8 (N=9/5). Dots represent biological replicates. Bars represent mean ± SEM. ***p<0.001, ****p<0.0001. (T) Top GO:BP terms following gene ontology analysis of differentially expressed genes in P7 MBNL1 cOE ventricles. (U) Heatmap showing expression of select cell cycle regulatory genes by RNAseq (N=3/4). See also Figure S5 and Figure S6.

### MBNL1 inhibits cardiomyocyte proliferation by stabilizing multiple cell cycle inhibitor transcripts

Comparisons of reciprocally regulated genes between MBNL1 gain and loss of function models identified cell cycle regulation among the top biological processes enriched in these gene sets (**Fig 6A-B**). Indeed, MBNL1 iKO ventricles had increased expression of positive cell cycle regulators and decreased expression of negative cell cycle regulators by RNAseq, which is the converse of the transcriptional phenotype observed in MBNL1 cOE ventricles (**Fig 6C**). Transcriptionally, positive regulators of proliferation appear to spike 7 days after cardiomyocyte-specific MBNL1 excision and remain elevated for another week relative to controls (**Fig S7A**). These gene expression changes were confirmed in isolated cardiomyocytes by qPCR (**Fig S7B-C**). Additionally, increased cell-cycle entry was confirmed histologically by increased numbers of cardiomyocytes which incorporated EdU and increased pH3 positivity (**Fig 6D-G**).

**Figure 6.**
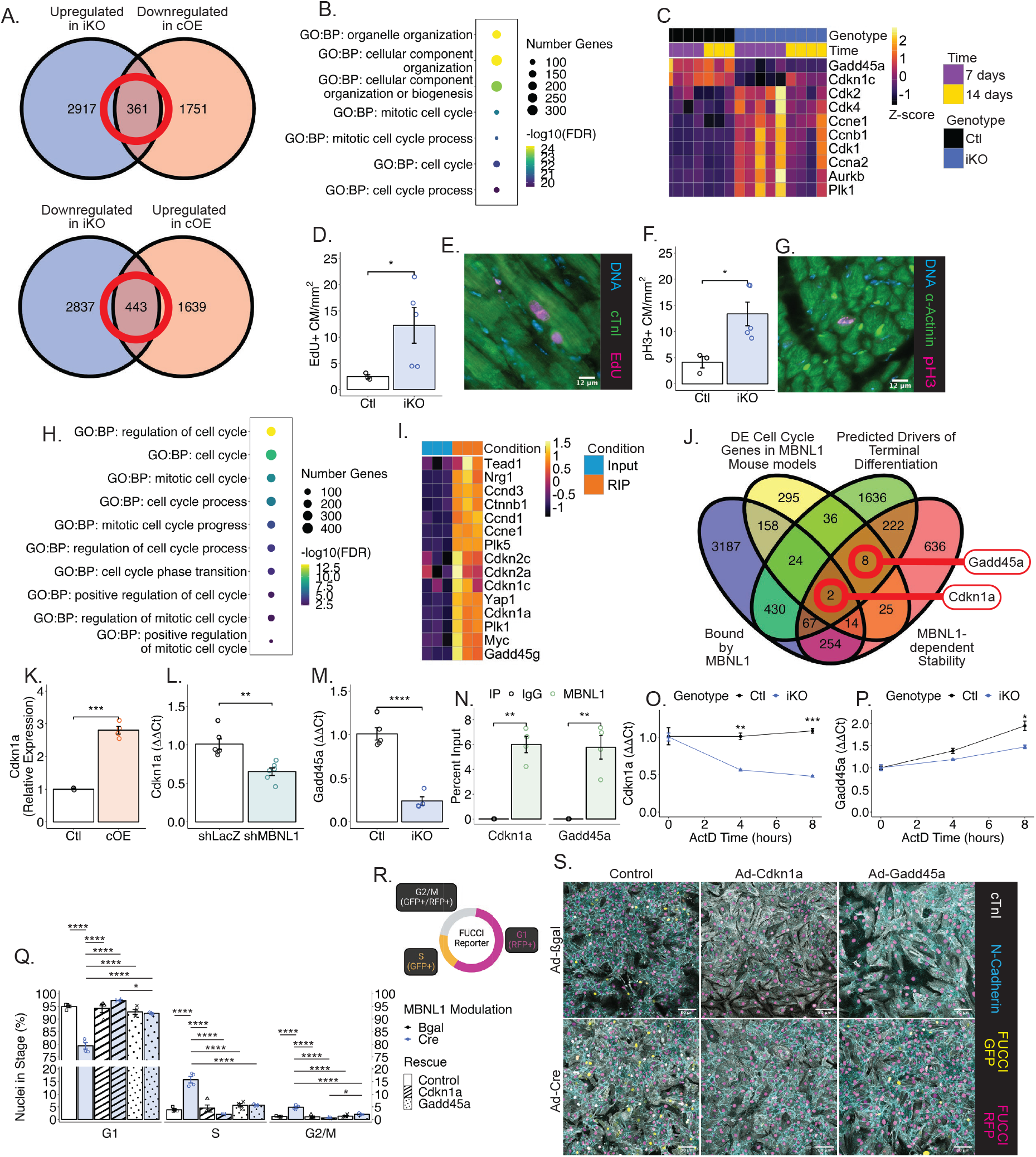
MBNL1 inhibits cardiomyocyte proliferation by stabilizing multiple cell cycle inhibitor transcripts. (A) Venn diagrams comparing reciprocally regulated genes between the MBNL1 iKO and MBNL1 cOE models. (B) Top GO:BP terms from gene ontology analysis of reciprocally regulated genes between MBNL1 iKO and MBNL1 cOE models. (C) Heatmap of select differentially expressed cell cycle regulatory genes by RNAseq in control versus MBNL1 iKO ventricles (N=7/9). (D-G) Quantification and representative images of EdU (D,E) and pH3 (F,G) positive cardiomyocytes (CM) by immunohistochemistry in control and MBNL1 iKO ventricles. Dots represent biological replicates. Bars represent mean ± SEM. *p<0.05. (H) Cell cycle relevant GO:BP terms from transcripts enriched in MBNL1 RIPseq. (I) Heatmap showing relative enrichment of cell cycle regulatory transcripts in MBNL1 RIPseq (N=3/3). (J) Venn diagram comparing cell cycle genes that were differentially expressed in at least one MBNL1 mouse model, genes that are bound by MBNL1, genes whose expression clustered with MBNL1 in our predicted drivers of terminal differentiation cluster, and genes that have MBNL1-dependent stability. (K) Expression of Cdkn1a in P7 control and MBNL1 cOE ventricles by RNAseq (N=3/4). Dots represent biological replicates. Bars represent mean ± SEM. ***p<0.001. (L) Expression of Cdkn1a by qPCR in adult rat ventricular myocytes treated with non-targeting (shLacZ) or MBNL1-targeted (shMBNL1) shRNA (N=6/6). Dots represent biological replicates. Bars represent mean ± SEM. **p<0.01. (M) Expression of Gadd45a by qPCR in isolated control and MBNL1 iKO cardiomyocytes (N=5/4). Dots represent biological replicates. Bars represent mean ± SEM. ****p<0.0001. (N) RIP-qPCR of Cdkn1a and Gadd45a transcripts in IgG versus MBNL1 RIP (N=4/4). Dots represent biological replicates. Bars represent mean ± SEM. **p<0.01. (O-P) Quantification of Cdkn1a (O) and Gadd45a (P) expression by qPCR following ActD treatment (N=3/3). Dots represent mean ± SEM. *p<0.05, **p<0.01, ***p<0.001. (Q-S) Quantification (Q), FUCCI color schematic (R), and representative images (S) of neonatal cardiomyocyte proliferation following adenoviral transduction using the FUCCI cell cycle reporter system (N=4). Dots represent biological replicates. Bars represent mean ± SEM. *p<0.05, ****p<0.0001. See also Figure S7.

To determine how MBNL1 could be regulating proliferation, the MBNL1 RIPseq dataset was mined for cell cycle regulators. Here, MBNL1-bound transcripts were significantly enriched for transcripts involved in regulation of cell cycle progression by gene ontology analysis (**Fig 6H**). In total, MBNL1 bound 266 transcripts with known function regulating the cell cycle, including previously identified central regulators of cardiomyocyte proliferation like Yap1/Tead1 and Myc as well as core proliferation machinery like cyclins, cyclin-dependent kinases, and cell cycle inhibitors (**Fig 6I**). This follows the developmental expression of MBNL1 which dramatically increases in expression at both the RNA and protein level just as cardiomyocytes begin to exit the cell cycle and hypertrophy becomes the predominant mechanism for cardiac growth^56^. To computationally identify likely transcripts by which MBNL1 could regulate cardiomyocyte proliferation, we compared cell cycle genes that were differentially expressed in at least one MBNL1 mouse model, genes that are bound by MBNL1, genes whose expression clustered with MBNL1 in our predicted drivers of terminal differentiation cluster, and genes that have MBNL1-dependent stability (**Fig 6J**). This analysis produced several likely regulatory targets including the cell cycle inhibitor Cdkn1a (encoding for p21) which we had previously identified as a MBNL1-dependent regulator of fibroblast proliferation^37^. Gadd45a was another cell cycle inhibitor that was identified as a transcript that satisfied all but the MBNL1 bound criteria. We chose to investigate this factor as Gadd45a was not identified in our RIPseq in this study due to one outlying sample but it was previously found to be regulated by MBNL1 in cardiac and embryonic fibroblasts^39^. Interestingly, both cell cycle inhibitors rise in expression over the same time course as MBNL1 (**Fig 1M, S7D-E**). Cdkn1a expression was strongly upregulated in MBNL1 cOE mice at all timepoints examined (**Fig 6K**). Additionally, Cdkn1a expression was decreased in adult rat ventricular cardiomyocytes following MBNL1 knockdown (**Fig 6L**) and Gadd45a expression was strongly downregulated in MBNL1 iKO myocytes (**Fig 6M**). To further confirm MBNL1-dependent regulation of these transcripts, MBNL1 was knocked down in rat NVMs. Here, MBNL1 knockdown decreased both Cdkn1a and Gadd45a expression (**Fig S7F-H**), which was recapitulated at the protein level for Gadd45a in MBNL1^F/F^ mouse NVMs following adenoviral Cre (AdCre) transduction (**Fig. S7I-J**). Furthermore, direct MBNL1 binding of both these cell cycle inhibitor transcripts was validated by RIP-qPCR, and a cluster of MBNL1 binding sites have been identified in the 3’UTR of the Cdkn1a transcript in previous MBNL1 CLIPseq studies (**Fig 6N, S7K**)^50^. qPCR validated that MBNL1 is essential for the stabilization of both Cdkn1a and Gadd45a (**Fig 6O-P**).

To directly determine whether MBNL1 suppresses proliferation by stabilizing Cdkn1a and Gadd45a transcripts, a rescue experiment was performed using the Fluorescent Ubiquitination-based Cell Cycle Indicator (FUCCI) biosensor (**Fig 6Q-S**). As expected, Cre-treated MBNL1^F/F^ mouse NVMs had significantly increased cell cycle activity compared to βgal-treated control NVMs, with approximately 4-fold increases in the number of myocytes in S- or G2/M-phases (**Fig 6Q**,**S**). Notably, reintroduction of Cdkn1a or Gadd45a blocked the increased cell cycle activity seen in MBNL1 KO NVMs (**Fig 6Q**,**S**). As Gadd45a had not been previously identified as a regulator of cardiomyocyte proliferation, we also performed a rescue experiment in the context of MBNL1 knockdown in NVMs. Here, MBNL1 knockdown resulted in a greater percentage Ki67-positivity in NVMs while overexpression of Gadd45a reduced this increased proliferation (**Fig S7L-M**). Together, this evidence suggests that MBNL1 acts to establish and maintain cardiomyocyte cell cycle exit at least in part by stabilizing cell cycle inhibitor transcripts Cdkn1a and Gadd45a.

### MBNL1 dosage controls the postnatal regenerative window

Cardiomyocyte differentiated cell state is integrally linked to cardiac regenerative potential^6^. Hence, it was hypothesized that the heart’s regenerative window could be tuned by modulating cardiomyocyte-specific MBNL1 activity. First, to determine if MBNL1 overexpression would decrease regeneration following injury in neonatal mice, left ventricular apical resection was performed on MBNL1 cOE and control littermates at P1 and hearts were harvested 7 and 21 days later (**Fig 7A**). At 7 days after injury all hearts independent of genotype showed apical scarring in picro sirius red stained myocardial sections (**Fig 7B-C**). By contrast, at 21 days after injury, when the heart is almost fully regenerated in control animals, MBNL1 cOE hearts had persistent apical scarring (**Fig 7B**,**D**) and poor systolic function (**Fig 7E**), which is consistent with impaired myocardial regeneration. Notably, apical resection control mice show no difference in systolic function relative to sham-operated age-matched controls, validating that wildtype mice fully regenerate after injury (**Fig 7E**). Furthermore, sham-operated MBNL1 cOE mice show no difference in function relative to sham-operated control animals at this age, suggesting that MBNL1 cOE hearts have functionally compensated for reduced cardiomyocyte proliferation at this timepoint, highlighting the regenerative perturbation as the driver for dysfunction in this model (**Fig 7E**). Following apical resection MBNL1 cOE mice also showed decreased ventricle weight/body weight ratios and increased cardiomyocyte cross-sectional areas, both consistent with impaired cardiac regeneration (**Fig 7F-G**). To further confirm that cardiomyocyte proliferation was inhibited in MBNL1 cOE mice, cardiomyocyte lineage-traced hearts were harvested 7 days after injury during the peak proliferative window. Significantly fewer MBNL1 cOE cardiomyocytes incorporated EdU relative to control mice, indicating that fewer cells had entered the cell cycle following injury (**Fig 7H-I**). Additionally, MBNL1 cOE mice had lower levels of Ki67-positivity (total proliferating cardiomyocytes) and pH3-positivity (mitotic cardiomyocytes) 7 days after injury when compared to controls (**Fig 7J-M**). Together, this evidence suggests that early MBNL1 overexpression is sufficient to prematurely close the postnatal regenerative window through its suppression of cell cycle activity.

**Figure 7.**
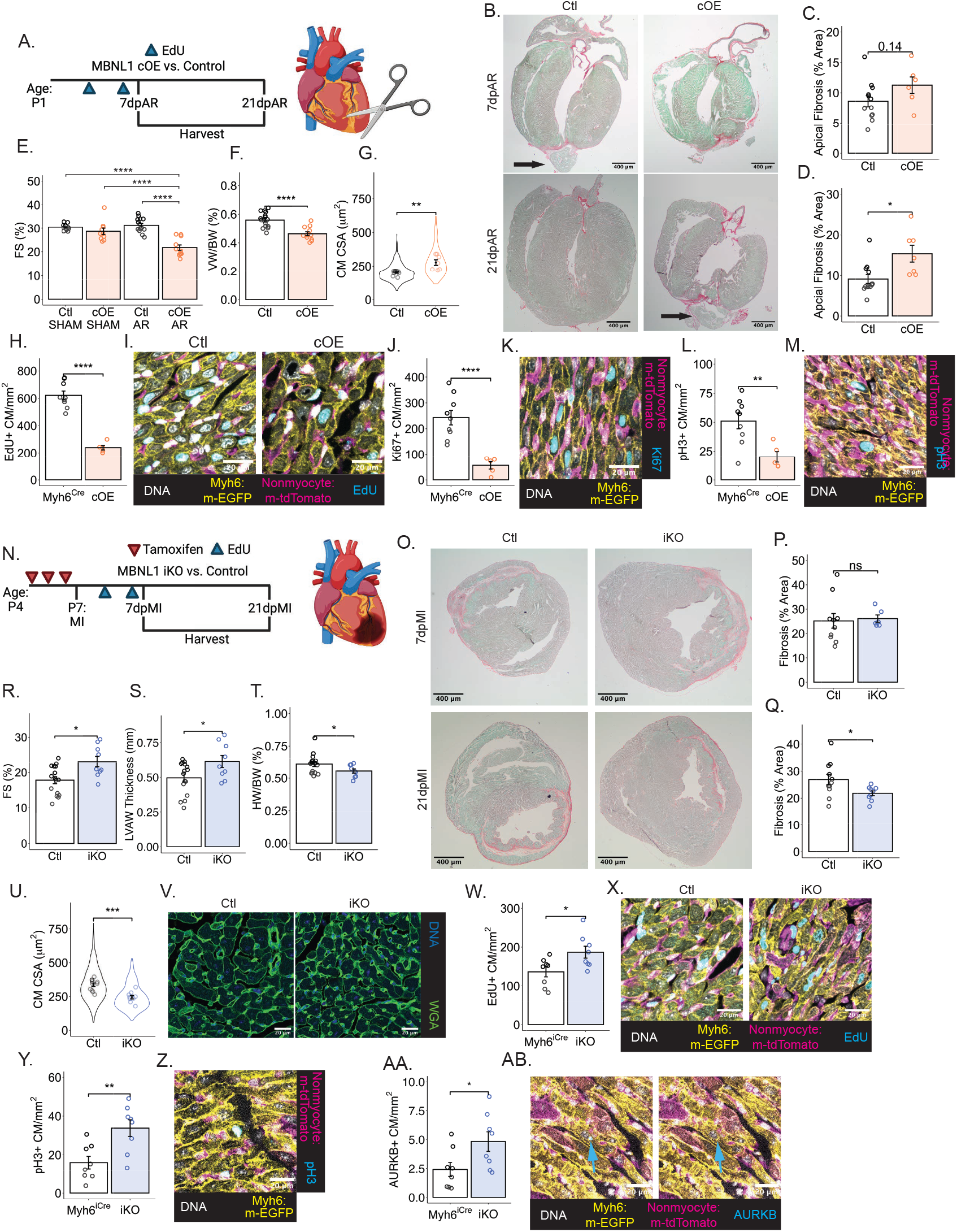
MBNL1 antagonizes cardiac regeneration. (A) Experimental schematic for P1 apical resection in MBNL1 cOE versus control mice. (B) Representative images of sirius red/fast green-stained myocardial sections 7 days (7dpAR, top) and 21 days (21dpAR, bottom) following apical resection. (C-D) Quantification of apical fibrosis shown in B at 7d (C, N=12/6) and 21d (D, N=12/7) following apical resection. Dots represent biological replicates. Bars represent mean ± SEM. *p<0.05. (E) Quantification of fractional shortening (FS) by echocardiography (N=11-14). Dots represent biological replicates. Bars represent mean ± SEM. ****p<0.0001. (F) Quantification of ventricle weight/body weight (VW/BW) ratio (N=16/12). Dots represent biological replicates. Bars represent mean ± SEM. ****p<0.0001 (G) Quantification of cardiomyocyte (CM) cross sectional area (CSA) in myocardial sections stained with WGA 21dpAR (N=11/7). Dots represent average CSA for an individual mouse and error bars represent the mean ± SEM. Violin plots represent the distribution of individual myocytes. Mouse average CSA was compared between groups. **p<0.01. (H-M) Quantification and representative images of EdU (H-I), Ki67 (J-K), and pH3 (L-M) positive cardiomyocytes (CM) 7d post apical resection (N=9/5). Dots represent biological replicates. Bars represent mean ± SEM. **p<0.01, ****p<0.0001. (N) Experimental schematic for P7 neonatal myocardial infarction (MI) in MBNL1 iKO versus control mice. (O) Representative images for sirius red/fast green-stained myocardial sections 7 days (7dpMI, top) and 21 days (21dpMI, bottom) following MI. (P-Q) Quantification of fibrosis shown in O at 7d (C, N=10/6) and 21d (D, N=11/9) following MI. Dots represent biological replicates. Bars represent mean ± SEM. *p<0.05, ns=not significant. (R-S) Quantification of fractional shortening (FS, R) and left ventricular anterior wall (LVAW) diastolic thickness (S) by echocardiography (N=16/9). Dots represent biological replicates. Bars represent mean ± SEM. *p<0.05. (T) Quantification of heart weight/body weight (HW/BW) ratio (N=16/9). Dots represent biological replicates. Bars represent mean ± SEM. *p<0.05. (U-V) Quantification (U) and representative images (V) of cardiomyocyte (CM) cross sectional area (CSA) by WGA staining 21dpMI. Dots represent average CSA for an individual mouse and error bars represent the mean ± SEM. Violin plots represent the distribution of individual myocytes. Mouse average CSA was compared between groups. ***p<0.001. (W-AB) Quantification and representative images of EdU (W-X), pH3 (Y-Z), and AURKB (AA-AB) positive cardiomyocytes (CM) 7days following myocardial infarction (N=8/8). Dots represent biological replicates. Bars represent mean ± SEM. *p<0.05, **p<0.01.

To determine if disrupting cardiomyocyte terminal differentiation via MBNL1 loss of function could be used to promote cardiac regeneration, MBNL1 iKO and control littermates were subjected to myocardial infarction (MI) via permanent left anterior descending artery (LAD) occlusion at postnatal day 7 (**Fig 7N**), which is at the end of the heart’s regenerative window in mice^58^. Here the MI model was used due to higher mortality of the apical resection model at later times during postnatal development. Following neonatal MI, all hearts independent of genotype showed left ventricular scarring along the LAD distribution 1 week after injury by picro sirius red staining (**Fig 7O-P**). By contrast, 21 days after injury MBNL1 iKO mice had significantly less fibrosis than their wildtype littermates (**Fig 7O**,**Q**). Additionally, MBNL1 iKO mice had significantly improved systolic function and thicker left ventricular anterior walls compared to control mice by echocardiography (**Fig 7R-S**). MBNL1 iKO mice had decreased heart weight/body weight ratios compared to control mice, as well as decreased cardiomyocyte cross sectional area 21 days following injury (**Fig 7T-V**). To determine if greater cardiomyocyte proliferation caused the heightened regeneration and smaller overall hearts following MI, cardiomyocytes were lineage-traced and myocardial sections examined for myocyte-specific EdU incorporation at 7 days after injury. Here, MBNL1 iKO cardiomyocytes had significantly increased EdU incorporation compared to control cardiomyocytes, indicating that more cardiomyocytes re-entered the cell cycle following injury (**Fig 7W-X**). MBNL1 iKO mice also had higher levels of pH3-positivity (mitotic cardiomyocytes) and AURKB-positivity (cytokinetic cardiomyocytes) indicating that there is more cardiomyocyte mitosis and true cardiomyogenesis occurring in MBNL1 iKO mice relative to controls (**Fig 7Y-AB**). Together, these data demonstrate that loss of MBNL1 function extends the neonatal cardiac regeneration window and enhances the heart’s regenerative capacity.

## Discussion

This study demonstrates that MBNL1 acts as a switch between immature/regenerative and terminally differentiated cardiomyocyte transcriptomes and the requirement for its activity in maintaining differentiated cardiomyocyte states in fully developed mice. MBNL1’s stabilization of transcripts that encode for critical promoters of cardiomyocyte maturity like the ERRα^44,45^ and all its downstream signaling targets (**Fig 4**) appears to be a dominant mechanism versus its other modes of modulating RNA metabolism (**Fig 3**). Indeed, RNA regulon theory posits that groups of transcripts encoding proteins in the same functional pathways are coordinately regulated via multiple levels of RNA metabolism to quickly activate entire pathways^59,60^, demonstrating MBNL1’s powerful transcriptome-wide function that turns fetal regeneration programs off while simultaneously turning on terminally differentiated states (**Fig 1**).

Importantly, this sheds new light on the role of post-transcriptional regulatory elements as regulators of cardiomyocyte state space. This fits into a growing body of work which has observed that differentiated cardiomyocyte states are a critical aspect governing cardiomyocyte regeneration (**Fig 5, 6**) in that cardiomyocytes must transition to a more fetal-like state to clonally expand^2–6^. Although past attempts at promoting cardiac regeneration through forced dedifferentiation of cardiomyocytes have been successful, perhaps it is not necessary to wipe away the cell’s entire history with Yamanaka factor reprogramming to achieve this goal^15^. These findings indicate that modulation of endogenous trans-acting factors controlling terminal differentiation programs is sufficient to tweak the developmental state of the heart thereby extending its regenerative window and capacity (**Fig 7**). Such knowledge is vital to leveraging controllers of cardiomyocyte state trajectories for therapeutics without causing off-target effects or driving the system too far down a dedifferentiation program to undesirable neoplasms^15^. Additionally, more work is needed to determine the causal relationship between cardiomyocyte dedifferentiation and proliferation. Although dedifferentiation seems to be a necessary precursor to cardiomyocyte proliferation, it does not appear that dedifferentiation alone in the absence of pro-proliferative signals is sufficient to drive regeneration^15^. In all likelihood, a more targeted combined approach relying on cardiomyocyte-specific developmental factors for (1) controlled dedifferentiation and (2) constitutive pro-proliferative signaling will be necessary for improving the efficacy of these approaches for eventual translation to the clinic.

Lastly, these results emphasize that if restoring the regenerative window in the adult heart is to be targeted clinically, a cautious and rationally guided approach is necessary. Prolonged loss of a cardiomyocyte’s mature cell state has created functional depression in multiple models of suppressed differentiation, signifying that such approaches must be further refined and tightly controlled (**Fig 2**)^9,11,13,15^. Additionally, we have previously shown that adult global MBNL1 knockout mice have a much higher risk of cardiac rupture and greatly suppressed cardiac function following MI^39^, meaning that calibrating the timing of therapeutic interventions to avoid exacerbation of known post-injury sequelae that could be adversely affected by loss of differentiation will likely be critically important in this realm.

## Acknowledgements

We would like to thank Drs. Rong Tian, Stephen Tapscott, Jeffery Chamberlain, and Joelle Chamberlain for their insight and helpful discussions of this work. Additionally, we would like to thank Dr. Dale Hailey for assistance with light microscopy and Edward Parker for assistance with electron microscopy. We would like to thank the following investigators for use of their plasmids deposited on Addgene: tFucci(CA)5 was a gift from Atsushi Miyawaki (Addgene plasmid # 153521; http://n2t.net/addgene:153521; RRID:Addgene_153521), U6 pShuttle was a gift from Ronald Kahn (Addgene plasmid # 13428; http://n2t.net/addgene:13428; RRID:Addgene_13428), pSIL-eGFP was a gift from Johannes Hell (Addgene plasmid # 52675; http://n2t.net/addgene:52675; RRID:Addgene_52675), MCS-BioID2-HA was a gift from Kyle Roux (Addgene plasmid # 74224; http://n2t.net/addgene:74224; RRID:Addgene_74224). Summary/schematic figures (**Fig 1A, 2A, 2N, S2B, 3A, 3D, 3I, 5A, 5M, S6A, 6R, 7A, 7N**) were created with Biorender. This work was supported by the National Institutes of Health (R01HL141187, R01HL142624, and R01HL162229 to JD; P30AR074990 and RM1GM131981 to MR; R01HL058493 to DPK; P30AG013280 to MJM; F30HL163926 to LRJB) and American Heart Association (Predoctoral Fellowship #905060 to LRJB).

## Author Contributions

LRJB, DB, IMR, CDO, JG, RJ, MJM, DPK, MR, and JMD conducted experiments and analyzed results. Experiments were designed by LRJB and JD. LRJB and JD contributed to writing and reviewing the manuscript.

## Declaration of Interests

The authors declare no competing interests.

## Materials and Methods

### Animal models

MBNL1 Tg^39^, MBNL1^F/F 37^, Myh6^iCre^ (JAX stock #005650)^46^, Myh6^Cre^ (JAX stock #011038)^52^, and mTmG (JAX stock #007576)^57^ mice have all been previously described. Cardiomyocyte-specific tamoxifen-inducible MBNL1 loss of function mice were generated by crossing MBNL1^F/F^ mice with Myh6^iCre^ mice to produce MBNL1^F/F^ Myh6^iCre^ (MBNL1 iKO); MBNL1^F/F^ and Myh6^iCre^ mice were used as controls. Cardiomyocyte-specific MBNL1 overexpression mice were generated by crossing MBNL1 Tg mice with Myh6^Cre^ mice to produce MBNL1 Tg Myh6^Cre^ (MBNL1 cOE); non-transgenic and Myh6^Cre^ mice were used as controls. Cardiomyocyte-specific tamoxifen-inducible MBNL1 overexpression mice were generated by crossing MBNL1 Tg mice with Myh6^iCre^ mice to produce MBNL1 Tg Myh6^iCre^ (MBNL1 iOE); non-transgenic and Myh6^iCre^ mice were used as controls. Lineage-traced cardiomyocytes were used for quantifying cardiomyocyte proliferation in neonatal regeneration models. Here, mTmG mice were crossed with MBNL1 cOE and Myh6^Cre^ control mice or MBNL1 iKO and Myh6^iCre^ control mice. All cells in mTmG mice express a membrane-targeted tdTomato (mT) unless Cre activity excises the mT and moves a membrane-targeted EGFP (mG) sequence in frame for expression. This allows for simultaneous labeling of cardiomyocytes (mG+) and non-cardiomyocytes (mT+). For tamoxifen inducible models, strategies for tamoxifen induction of Cre activity are presented in relevant figures; all mice in these studies received tamoxifen. Here, mice received intraperitoneal (IP) injections of 25mg/kg (adult) or 12.5mg/kg (neonate) pharmaceutical grade tamoxifen dissolved in 95% peanut oil and 5% ethanol. In Figure 2C-M, tamoxifen citrate chow (400mg/kg chow, Harlan Laboratories) was given to mice after tamoxifen injections were completed to ensure full knockout of MBNL1. Wild type Sprague Dawley rats were used for experiments utilizing rat cardiomyocytes. Experimenters remained blinded to the genotypes until analysis was complete. Both male and female animals were used in all experiments and mice were randomly assigned to groups. Echocardiography was performed on a Vevo2100 or Vevo3100 platform under light isoflurane anesthetic. All animal experiments were reviewed and approved by the University of Washington Institutional Animal Care and Use Committee (IACUC).

### Apical resection and myocardial infarction models

Neonatal apical resection and myocardial infarction models were performed as previously described^61^. Briefly, all P1 or P7 mice were separated from their parents. One at a time, neonatal mice were anesthetized using hypothermia. Mice were then immobilized on a pre-cooled platform and a lateral thoracotomy was performed to expose the left ventricle. For apical resection, the heart was then exteriorized by applying pressure to the abdomen and iridectomy scissors were used to remove approximately 15% of the left ventricle at the apex. Left ventricular chamber exposition was used as an anatomical landmark to ensure uniform injury between mice. For MI, the left anterior descending artery (LAD) was permanently ligated approximately 1mm below the left atrium using an 8-0 Surgipro tapered suture. LAD ligation was confirmed by visualization of LV blanching below the ligation. Finally, the lateral thoracotomy and skin incisions were sutured closed, and the mouse was rapidly re-warmed before being returned to its litter. Once all surgeries were complete, the litter was reintroduced to their parents. Experimenters were blind to genotype at the time of surgery and remained blinded to genotypes until all measurements were taken and analyses were completed. For *in vivo* EdU pulse-chase experiments, EdU dissolved in PBS (100mg/kg) was administered by intraperitoneal injection 3- and 6-days after injury before harvesting at 7-days after injury.

### Cardiomyocyte isolation and cell culture

Mouse ventricular myocytes were freshly isolated by Langendorff perfusion with Liberase TM (0.225 mg/mL, Roche) in Krebs-Henseleit buffer (135mM NaCl, 4.7mM KCl, 0.6mM KH_2_PO_4_, 0.6mM Na_2_HPO_4_, 1.2mM MgSO_4_, 20mM Hepes,10μM BDM, and 30mM Taurine) as previously described^62^. Ventricular myocytes were mechanically dispersed and filtered through a 200μm nylon mesh then allowed to sediment for 5-10 minutes. Sedimentation was repeated three times using increasing Ca^2+^ concentration from 0.125 to 0.25 to 0.5 mmol/L. Myocytes were then harvested for downstream applications or plated on laminin-coated coverslips in Tyrodes solution (137mM NaCl, 5.4mM KCl, 0.5mM MgCl_2_, 1.2mM CaCl_2_^*^2H_2_O, 10mM HEPES, and 5mM Glucose, 7.4 pH) for 1 hour at 37°C prior to functional measurements. For myocyte morphology measurements, myocytes were relaxed in Tyrodes solution containing 25μM blebbistatin for 1 hour at 37°C and subsequently fixed with 4% PFA at room temperature for 15 minutes.

Adult rat ventricular cardiomyocytes were freshly isolated by Langendorff perfusion with Liberase TM (0.225 mg/mL, Roche) in modified Krebs-Henseleit buffer (135mM NaCl, 5.31 mM KCl, 0.33mM NaH_2_PO_4_, 1mM MgCl_2_, 20mM HEPES, 5.5mM glucose, 2mM sodium pyruvate, 10mM taurine, and 10mM BDM). Ventricular myocytes were mechanically dispersed and filtered through a 200μm nylon mesh then allowed to sediment for 5-10 minutes. Sedimentation was repeated four times using increasing Ca^2+^ concentration from 0.25 to 0.5 to 0.75 to 1mM. Myocytes were then resuspended in culture media (M199 supplemented with 1x insulin-transferrin-selenium, 1x penicillin-streptomycin, 5mM taurine, 1mM sodium pyruvate, 5mM creatine, and 2mM L-carnitine) and plated on laminin-coated wells to attach for 2 hours. After 2 hours, the culture media was replaced with a minimal volume of adenovirus-containing culture media at a multiplicity of infection (MOI) of 1000. One hour later, the adenovirus-containing media was diluted with fresh culture media to adequately fill the culture vessel. Following overnight culture, the media was replaced with fresh culture media. Cells were harvested 5 days after transduction.

Neonatal murine ventricular cardiomyocytes were freshly isolated from mouse or rat ventricular tissue at P1-2. Ventricles were enzymatically dissociated in ADS buffer (116.3mM NaCl, 19.9mM HEPES, 10.8mM NaH2PO_4_, 5.5mM glucose, 5.4mM KCl, and 0.83mM MgSO_4_) containing 5mg/mL Type II Collagenase, 1mg/mL Pancreatine, and 5U/mL DNAseI. Dissociated cells were passed through a 70μm filter and pre-plated on uncoated tissue culture plastic for at least 1.5 hours at 37°C to deplete non-myocytes. Ventricular myocytes were then collected, pelleted, resuspended, and plated on gelatin-coated tissue culture dishes at a density of 1.25*10^5^ cells/cm^2^ (rat) or 1.5*10^5^ cells/cm^2^ (mouse). Cells were transduced with adenoviruses at the time of plating at an MOI of 200 unless otherwise noted. Plating media (67.2% DMEM, 16.8% M199, 10% horse serum, 5% fetal bovine serum, 1% penicillin/streptomycin) was used for the first 24 hours of culture after which cells were maintained in maintenance media (46.9% DMEM, 46.9% M199, 5% fetal bovine serum, 1% penicillin/streptomycin, 0.2% insulin-transferrin-selenium, 0.1% chemically defined lipid).

### Gene expression analysis and bioinformatics

Gene expression analysis methods are previously described from our laboratory^37,39,63,64^. Briefly, total RNA was extracted using TRIzol and the Qiagen RNeasy kit for *in vivo* samples or the Zymo Direct-zol RNA Microprep kit for *in vitro* cell culture samples. For quantitative polymerase chain reaction (qPCR), total RNA was reverse transcribed into cDNA using random hexamer primers and SuperScript III first-strand synthesis kit (Invitrogen) according to the manufacturer’s instructions. RT-PCR was performed on a CFX96 Real-Time System with a Biorad C1000 Touch Thermal Cycler using SsoAdvanced SYBR Green (Biorad). Thermocycler conditions were as follows: Polymerase Activation and DNA Denaturation at 95^°^C for 30s followed by 40 total cycles of Denaturation at 95°C for 10s and Annealing/Extension at 60°C for 30s followed by plate read. Gene expression fold change was determined using the delta-delta Ct method. Gapdh or Rps18 served as housekeeping genes. A list of primers used for qPCR can be found in Table S1.

For RNA sequencing (RNAseq), sequencing was performed by BGI Genomics using paired-end 100 nucleotide sequencing with 20-40 million reads depending on the experiment. Cleaned reads were aligned to reference genomes (mouse: GRCm38, rat: Rnor6.0) using STAR with de novo junction discovery and summarized at the gene level using featureCounts^65,66^. Differential expression analysis was completed using DESeq2 using the standard workflow^67^. For alternative splicing analysis, reads were quantified at the transcript level using Salmon and alternative splicing analysis was completed using IsoformSwitchAnalyzeR^68,69^. Pathway and ontology enrichment analyses for all experiments were performed using gprofiler2^70^. Heatmaps were generated using the pheatmap package in R. To generate gene-gene correlation data in **Fig 1**, differential expression analysis was performed between all timepoints, and gene-gene spearman correlations were calculated for all genes that were differentially expressed in at least one comparison. K-nearest neighbor (KNN) clustering was subsequently performed on the spearman correlations using 8 clusters as this was the number of large clusters revealed by hierarchical clustering and clustering with more than 8 clusters began to produce small clusters with few constituents.

### RNA immunoprecipitation in neonatal cardiomyocytes

Neonatal cardiomyocytes were isolated as stated above. Cells were adenovirally transduced with a FLAG-tagged MBNL1 construct (AdMBNL1-FL, MOI 100) for 48hours. Myocytes were scraped in cold 1X PBS and pelleted by centrifugation for 3 minutes at 1000rpm. Cells were then lysed in RIP buffer [150mM KCl, 25mM Tris-HCl pH 7.4, 5mM EDTA, 0.5% NP-40, 0.5mM DTT, 100U/mL RNAse OUT Recombinant Ribonuclear Inhibitor (ThermoFisher 10777019), and 1X complete EDTA-free Protease Inhibitor Cocktail (Sigma 11836170001)]. Lysates were mechanically sheared by Dounce homogenization with a type B (tight) pestle using 18 strokes. Nuclear membrane and debris were pelleted by centrifugation at 13000 rpm for 10 minutes at 4°C. For each IP, 10% of the input lysate was placed directly into TRIzol; lysates were then incubated overnight with 5μg of either Rabbit-anti-FLAG (Sigma F7425) or Rabbit IgG Isotype control (Cell Signaling 3900S) antibody at 4°C with gentle rotation. Following overnight incubation, 40μL of Dynabeads Protein A (ThermoFisher 10001D) were washed 6x in RIP buffer and then added to RIP lysates for 1 hour at 4°C with gentle rotation. Beads and bound material were collected using a DynaMag magnet, and the beads were washed 6x in RIP buffer and 1x in PBS containing 0.5mM DTT, 100U/mL RNAse OUT, and 1x complete EDTA-free Protease Inhibitor Cocktail. Following washing, TRIzol was added to isolate bound RNA. Following RNA isolation, RNA was quantified by RNA-seq (comparing input and IP RNA) or converted to cDNA and measured by qPCR using the percent input method.

### MBNL1 interactome labelling using the BioID system

#### Sample preparation

The BioID system was used as previously described for MBNL1 interactome labeling^48^. Neonatal rat ventricular cardiomyocytes were isolated as stated above. Cells were treated with AdBioID2-HA or AdMBNL1-BioID2-HA (MOI 100) overnight. 48 hours after transduction, cells were treated with 50μM biotin for 16 hours. Cells were then washed twice with PBS, scraped in cold PBS, pelleted, and lysed in lysis buffer (50mM Tris, pH 7.4, 500mM NaCl, 0.4% SDS, 2% Triton-X-100, 1mM DTT, 1x complete protease inhibitor [Roche]). Lysates were sonicated twice for 1 minute at 30% duty cycle at an output level of 4. Lysates were then diluted 1:1 in 50mM Tris pH 7.4 and centrifuged at 16,500g for 10 minutes at 4°C to pellet debris. Supernatants were then incubated overnight with 300ul of pre-washed streptavidin conjugated Dynabeads overnight at 4°C with gentle rotation. Beads were then connected with a magnetic stand and washed twice in 2% SDS, once in wash buffer 1 (0.1% deoxycholate, 1% Triton X-100, 500mM NaCl, 1mM EDTA, 50mM HEPES, pH 7.5), once in wash buffer 2 (250mM LiCl, 0.5% NP-40, 0.5% deoxycholate, 1mM EDTA, 10mM Tris, pH 8), and once in 50mM Tris, pH 7.4. 10% of the sample was saved for western blot and the remaining 90% of the sample was further processed for mass spectroscopy. The magnetic streptavidin beads were further washed three times with 200μL of 600mM NaCl and 1% Triton X100 for 10 minutes using a thermomixer operated at 2000 rpm and 37°C. The salt and detergent were removed by briefly washing three more times with 50 mM Tris pH 8 at room temperature. The disulfide bonds of any proteins attached to the beads were reduced in 50μl 0.1% PPS Silent Surfactant (no longer available), 50mM Tris pH 8, and 10mM tris(2-carboxyethyl)phosphine (TCEP) for an hour in a thermomixer at 47°C and 2000 rpm. Cysteine thiols were alkylated by the addition of 11mM iodoacetamide at room temperature for 20 minutes. The beads were then mixed with 0.5μg sequencing grade trypsin (Pierce), and digestion proceeded for five hours at 37°C. The resulting supernatants were removed and placed in new vials. The beads were washed briefly using 50μL 0.1% trifluoracetic acid in water, and these washes were combined with the supernatants in a final volume of ∼100μL per sample. Negative control samples were run prior to the BioID samples eliminating the possibility of carryover contamination of the former.

#### Liquid chromatography-mass spectrometry

All mass spectrometry was performed on a Q-Exactive HF (Thermo Fisher Scientific) mass spectrometer with a Waters Nanoacquity HPLC. The Nanoacqity autosampler was used to inject 7μL of each tryptic digest onto a 2 cm x 150-μm Kasil fritted trap (Dr. Maisch Reprosil-Pur 120 C18-AQ 3 μm beads) at a flow rate of 2μL/min. After loading and desalting, the trap was brought on-line with a 100μm x 30 cm Kasil fritted column packed with the same Dr. Maisch beads. The outlet of the column was attached to an empty pulled tip (20μm ID) via a zero dead volume connector. The column and trap were mounted to a nanospray ion source (CorSolutions, Ithaca, NY) heated to 50°C, and placed in line with the HPLC pump. Peptides were eluted off the column using a gradient of 2-40% acetonitrile in 0.1% formic acid over 105 minutes, followed by 40-60% acetonitrile over an additional 15 minutes at a flow rate of 0.45μL/min. The mass spectrometer was operated using electrospray ionization (2 kV) with the heated transfer tube at 300°C, where each sample was analyzed using data dependent acquisition (DDA). For DDA, one orbitrap mass spectrum (m/z 400-1600) was acquired followed by 20 orbitrap MS/MS spectra. The resolution for MS in the orbitrap was 60,000 at m/z 200, and 15,000 for MS/MS. The automatic gain control targets for MS and MS/MS were 3e6 and 1e5, respectively. The maximum fill time was 25 msec. The quadrupole isolation width was 1.4 m/z and HCD collision energy was 27%. Furthermore, MS/MS acquisitions were allowed for precursor charge states of 2-4. Dynamic exclusion (including all isotope peaks) was set for 10 seconds using monoisotopic precursor selection with a mass error of 10 ppm. Database searches were performed using Comet, and post-processing was done using PeptideProphet and ProteinProphet. Peptide hits were subsequently summarized at the protein level using ProteinProphet ^71^. Differential protein enrichment was calculated using the limma package with spectral counts as input ^72^.

### Transcript stability assay

Adult mouse ventricular cardiomyocytes were isolated and plated on laminin-coated coverslips 7 days following tamoxifen induction. Following attachment, transcription was inhibited with 2μg/mL Actinomycin D (Sigma), and RNA was isolated at baseline and 4 and 8 hours later for analysis by RNA-seq or qPCR. For analysis of RNAseq data, raw counts were normalized to transcripts per million (TPM) and the fraction of RNA remaining at 4 and 8 hours was calculated for each transcript with an average of >10 TPM at baseline. To account for differences in sequencing depth and RNA composition, RNA fractions were normalized using an adaptation of DESeq2’s median of ratios method in which the geometric mean of the fraction of remaining RNA was calculated for each transcript and then sample-specific normalization factors were determined by calculating the median ratio of all ratios of the fraction of RNA remaining compared to the geometric mean of remaining RNA for each transcript^67^. Subsequently, a moderated t-test was implemented from the limma package to compare normalized fractions of RNA remaining between genotypes^72^.

### Measurements of myocyte contractility and calcium transients

Sarcomere measurements were obtained from isolated myocytes using the IonOptix™ SarcLen Sarcomere Length Acquisition Module with a MyoCam-S3 digital camera (Ionoptix Co., Milton, MA) attached to an Olympus uWD 40 inverted microscope. For these measurements myocytes were bathed in 1.2mM Ca^2+^ Tyrode’s buffer (137mM NaCl, 5.4mM KCl, 0.5mM MgCl_2_, 1.2mM CaCl_2_^*^2H_2_O, 10mM HEPES, 5mM Glucose, 7.4 pH) and kept at 37°C. Sarcomere lengths were then measured in real time at a frequency of 1Hz and averaged across 10-15 contraction cycles. Separate coverslips were treated with 1uM Fura-2–acetoxymethyl ester to measure calcium transients. Blinded analysis was performed using the IonWizard software. Statistical analyses were performed on individual myocyte measurements (n ∼ 20 myocytes/mouse; N=3-4) using a Student’s t-test.

### Western blot

Protein samples were lysed directly in concentrated Laemmli buffer (15% Glycerol, 1% SDS, 62.5mM Tris HCl, pH 6.8, 0.05% bromophenol blue, 60mM DTT, and 1x complete protease inhibitor). Lysates were further diluted in Laemmli buffer and sample concentrations were normalized by total protein (Coomassie) staining following SDS-PAGE electrophoresis. Equal amounts of protein were subsequently loaded into 10-15% SDS-PAGE gels and transferred to PVDF membrane for immunodetection. Endogenous peroxidase activity from mitochondrial proteins was quenched by incubating membranes in 3% H_2_O_2_ for 15 minutes following transfer. Membranes were blocked in 5% milk in tris buffered saline containing 0.1% Tween-20 (TBST), pH 7.6. Primary antibody was applied overnight at 4°C in blocking buffer (α-Actinin, Sigma A7811, 1:2500; AURKB, Sigma A5102, 1:1000; GADD45A, CST 4632, 1:1000; GAPDH, Fitzgerald 10R-2932, 1:10000; MBNL1, CST 94633, 1:1000; MBNL1,

Abcam ab45899, 1:1000; phospho-YAP S109, CST 53749, 1:1000; Total OXPHOS Rodent WB antibody cocktail, Abcam ab110413, 1:1000; Troponin I, Millipore MAB1691, 1:1000; YAP, CST 14074, 1:1000). HRP-conjugated secondary antibody (1:4000, Sigma) was applied the following day for 1 hour at room temperature. Total protein (Coomassie), GAPDH, and α-Actinin were used as loading controls depending on the experiment.

### Histology and immunohistochemistry

Mice were euthanized and hearts were fixed in 4% paraformaldehyde overnight. Tissues were processed through a sucrose gradient (5-30%), cut in half in on either the transverse or longitudinal plane, embedded in optimal cutting temperature compound (OCT), and prepared for 5μm cryosectioning. To quantify fibrosis, picro sirius red/fast green staining was used (0.1% Direct Red 80, 0.1% Fast Green in Picric Acid). This method stains muscle tissue in green and fibrotic scar in red. Images of whole hearts were taken at 2x magnification and quantified in ImageJ using color thresholding. Immunohistochemistry methods are previously described from our laboratory^37,63,64^. Briefly, OCT sections were rinsed twice in PBS and the blocked in a fish skin gelatin solution (PBS with 1% BSA, 0.1% cold fish skin gelatin, 0.5% Triton X-100, and 0.05% sodium azide, pH 7.2-7.4). For antibody-based staining, primary antibodies for Ki67 (1:200, abcam), phosphorylated histone H3 (Ser 10) (1:200, abcam), Aurora B Kinase (1:100, Sigma), alpha-actinin (1:100, Sigma), and cardiac troponin I (1:100, abcam) were incubated overnight in blocking solution at 4°C. For myocyte cross sectional area, myocytes were stained with Wheat Germ Agglutinin, Alexa Fluor 488 Conjugate (1:100, Invitrogen). Following 3 washes, samples were incubated in AlexaFluor secondary antibodies (1:1000, Invitrogen) for 1.5 hours at room temperature to detect the antigen. Hoechst 33258 (1:1000 Thermo Fisher) was used to visualize nuclei. To visualize EdU incorporation, Click chemistry was used following the manufacturer’s instructions (Click-iT™ EdU Cell Proliferation Kit, ThermoFisher). Wash steps were performed with PBS containing 0.5% Triton X-100. All samples were mounted using Mowiol 4-88.

### Immunocytochemistry

Immunostaining procedures followed previously described methods from our lab^37,63,64^. Briefly, cardiomyocytes were fixed in 4% paraformaldehyde for 15 minutes at room temperature, washed/permeabilized in 1X PBS containing 0.5% Triton X-100, and blocked in PBS containing 0.5% Triton X-100 and 5% Normal Goat Serum (NGS). For myocyte morphology tracings, myocytes were stained with Wheat Germ Agglutinin, Alexa Fluor™ 488 Conjugate (1:100, Invitrogen) for 1 hour at room temperature. For antibody-based staining, primary antibodies for α-actinin (Sigma, 1:400), cardiac troponin I (cTnI, abcam, 1:400), MBNL1 (Cell Signaling, 1:200), or N-Cadherin (Sigma, 1:200) were incubated overnight at 4°C. Cells were then washed in PBS and AlexaFluor secondary antibodies (1:1000, Invitrogen) were incubated on the sample for 1.5 hours at room temperature to detect the antigen. To detect biotin, an Alexa Fluor 488 streptavidin conjugate was used (1:500, Thermofisher). Following 2 washes in PBS, Hoechst 33258 (1:1000, Invitrogen) was applied for 10 minutes to visualize nuclei. Cells were washed once more in PBS and then mounted in Mowiol 4-88.

### Image analysis

To quantify cardiomyocyte proliferation markers in neonatal models, the mTmG mouse was used as described above to provide both a positive (mG+) and negative (mT+) marker for cardiomyocytes. Five sequential 1 μm z-stacks were acquired (the entire thickness of the section) so that signal could be viewed in 3 dimensions. To quantify EdU+ or Ki67+ cardiomyocytes *in vivo*, cells were manually counted across 6-10 representative fields of view (FOVs) from the border zone of 2 sections, and counts were normalized to tissue area. To score pH3+ or AURKB+ cardiomyocytes, cells were manually counted across large tissue stitches encompassing at least 1mm^2^ of tissue including the border zone and interventricular septum from 2 sections. To score EdU+ and pH3+ cardiomyocytes in adult MBNL1 iKO mice, cells were manually counted across the entirety of a transverse section at the level of the papillary muscle using α-actinin or cTnI to mark cardiomyocytes. To score cell cycle stages with the FUCCI cell cycle reporter, 49 representative fields of view (FOVs) were captured, and nuclei cell cycle stage was quantified using cell profiler and confirmed by manual scoring. For myocyte geometry quantifications, statistical analyses were performed on individual myocyte measurements (n ∼ 40 myocytes/mouse) using a Student’s t-test.

### Electron microscopy

Samples were immediately fixed in 4% Glutaraldehyde in 0.1M sodium cacodylate buffer, then stored overnight at 4°C. The tissue was then washed 5 times for 5 minutes in fixative buffer at room temperature and post-fixed in buffered 2% osmium tetroxide, on ice, for 1 hour. Five washes were performed in ddH_2_O, then samples were en-bloc stained in 1% Uranyl Acetate (aqueous) overnight at 4°C. The next day the tissue was washed 5 times for 5 minutes in ddH_2_O and then dehydrated in ice cold 30%, 50%, 70%, and 95% ethanol. Samples were then allowed to come to room temperature. This is followed by 2 changes of 100% ethanol and two changes of propylene oxide. The tissue was then infiltrated in a 1:1 mixture of propylene oxide:epon araldite resin for 2 hours followed by two changes of fresh epon araldite (2 hours each change). Samples were then placed in flat embedding molds and polymerized at 60°C overnight. Samples were subsequently sectioned at 80nm thickness and imaged at 80KV on a JEOL 1230 transmission electron microscope.

### Adenoviral gene transfer

MBNL1-FLAG, βgal, and Cre recombinase adenoviruses were previously described^39^. The human ERR-alpha and GFP adenoviruses were gifts from Dr. Daniel Kelly. Cdkn1a/p21 (Vector Biolabs #1041) and Gadd45a (Vector Biolabs # 1582) adenoviruses are commercially available. The BioID2 adenoviral vector was generated by cloning BioID2-HA (Addgene plasmid #74224) into the pShuttle-CMV vector. The MBNL1-BioID2 adenoviral vector was generated by cloning BioID2-HA onto the C-terminus of MBNL1 separated by a flexible linker region (MBNL1-SGGGGS-BioID2-HA) into the pShuttle-CMV vector. The FUCCI adenoviral vector was generated by cloning tFucci(CA)5 (Addgene plasmid #153521) into the pShuttle-CMV vector. For shRNA adenoviral vectors, a pShuttle-U6 vector (Addgene plasmid #13428) was modified by replacing the U6 promoter and multiple cloning site (MCS) with the U6 promoter, MCS, CMV promoter, and EGFP of pSIL-eGFP (Addgene plasmid #52675) to form a pShuttle-U6-MCS-CMV-GFP vector. shLacZ and shMBNL1 were subsequently cloned downstream of the U6 promoter. shRNA sequences are available in Table S2. All adenoviruses generated for this study were sequenced through the coding region to verify the correct coding sequence. pShuttle constructs were subsequently transfected into HEK293A cells using the AdEasy adenoviral production kit (Agilent Technologies). The recombinant adenovirus was plaque purified, expanded, and purified/concentrated by cesium chloride gradients.

### Statistical analysis and data visualization

R version 4.2.1 and ggplot2 version 3.3.6 were used for statistical analysis and plotting data. Data are represented as mean ± SEM unless otherwise stated. 1- or 2-way ANOVA with Tukey post hoc analysis was used for multiple comparisons depending on the experimental setup. Unpaired two-tailed t-tests were used for pairwise comparisons. p < 0.05 was considered significant. Significance is denoted on graphs in the following manner: * p<0.05, ** p<0.01, ***p<0.001, ****p<0.0001. Replicate numbers are given in figure legends; for pairwise comparisons numbers are given as N=control/experimental unless otherwise specified. For experiments with multiple groups, a range of replicate numbers is given unless otherwise specified.

## Supplementary Figure Legends

**Supplementary Figure 1.**
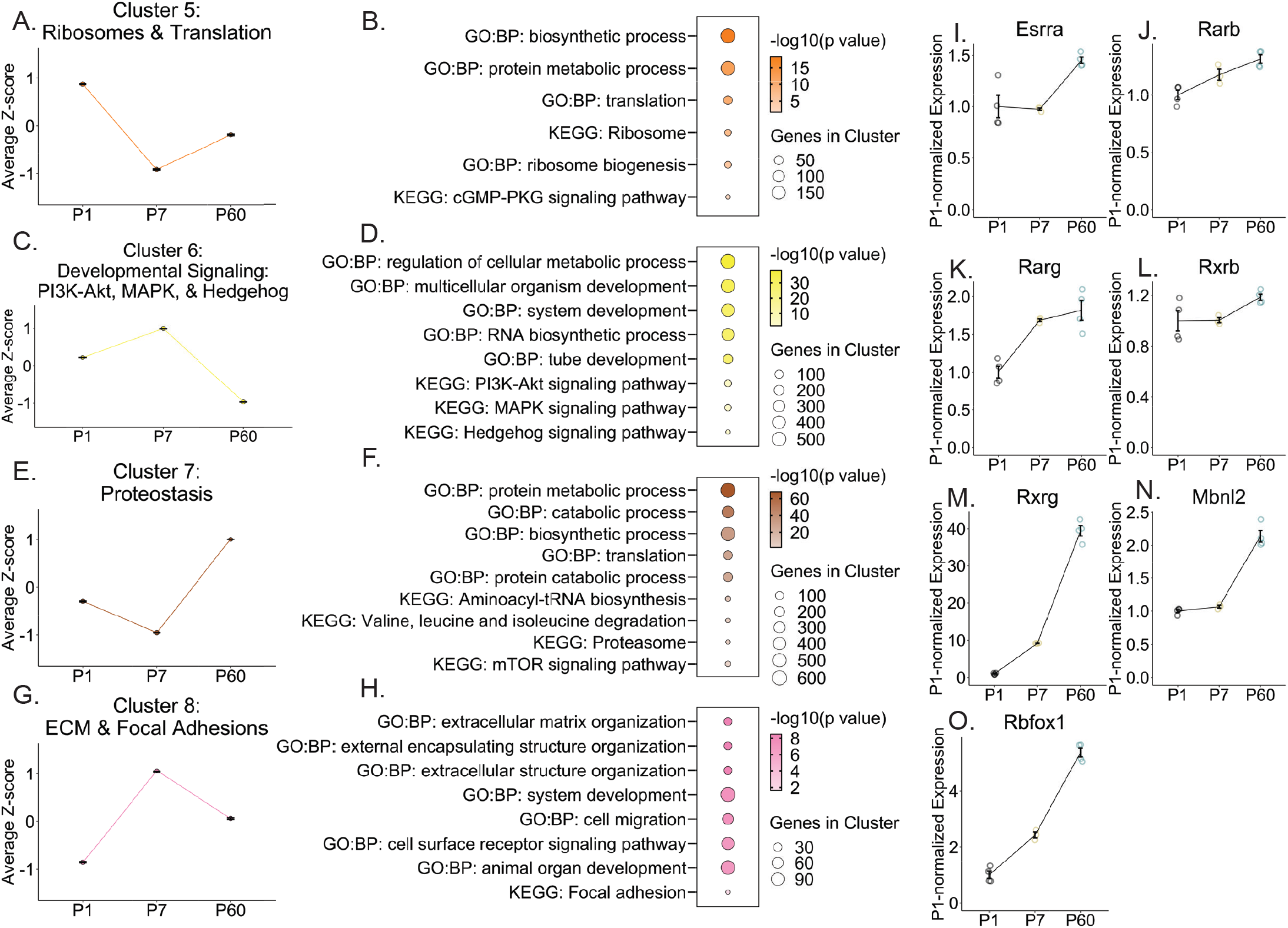
Further characterization of cardiomyocyte terminal differentiation, related to Figure 1. (A,C,E,G) Mean expression ± SEM of genes within each cluster over time in wild type ventricles. (B,D,F,H) Representative GO:BP & KEGG terms for each cluster. (I-O) Expression of Esrra (I), Rarb (J), Rarg (K), Rxrb (L), Rxrg (M), Mbnl2 (N), Rbfox1 (O) by RNAseq over time in wild type ventricles. Dots show biological replicates. Lines show mean ± SEM.

**Supplementary Figure 2.**
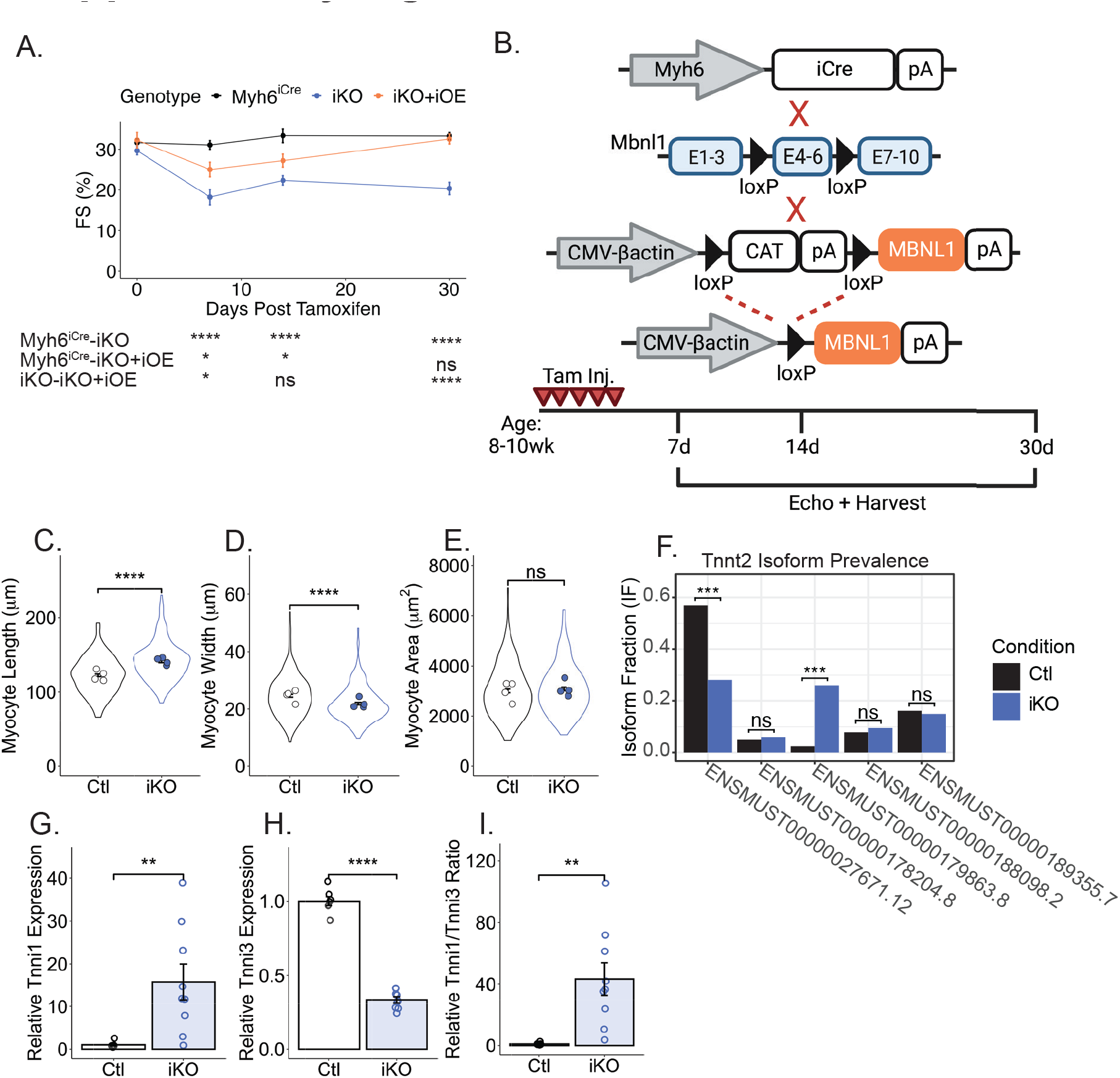
Further phenotypic characterization of MBNL1 iKO mice, related to Figure 2. (A) Quantification of left ventricular fractional shortening (FS) by echocardiography following tamoxifen induction (N=9-14). *p<0.05, ****p<0.0001. Dots represent mean ± SEM. (B) Breeding scheme to generate MBNL1 iKO+iOE mice (top) and experimental schematic (bottom). (C-E) Quantification of length (C), width (D), and area (E) of isolated control and MBNL1 iKO cardiomyocytes. Dots represent the mean for an individual mouse. Violin plots represent the distribution and error bars show the SEM of individual myocytes. Individual myocytes were compared between groups. N=200 myocytes/N=4 mice for each group. ****p<0.0001, ns=not significant. (F) Quantification of relative Tnnt2 isoform prevalence by RNAseq in control and MBNL1 iKO ventricles (N=7/9). Bars show mean. ***p<0.001, ns=not significant. (G-I) Expression of Tnni1 (G), Tnni3 (H), and ratio of Tnni1/Tnni3 (I) by RNAseq (N=7/9). Dots are biological replicates and bars are mean ± SEM. **p<0.01, ****p<0.0001

**Supplementary Figure 3.**
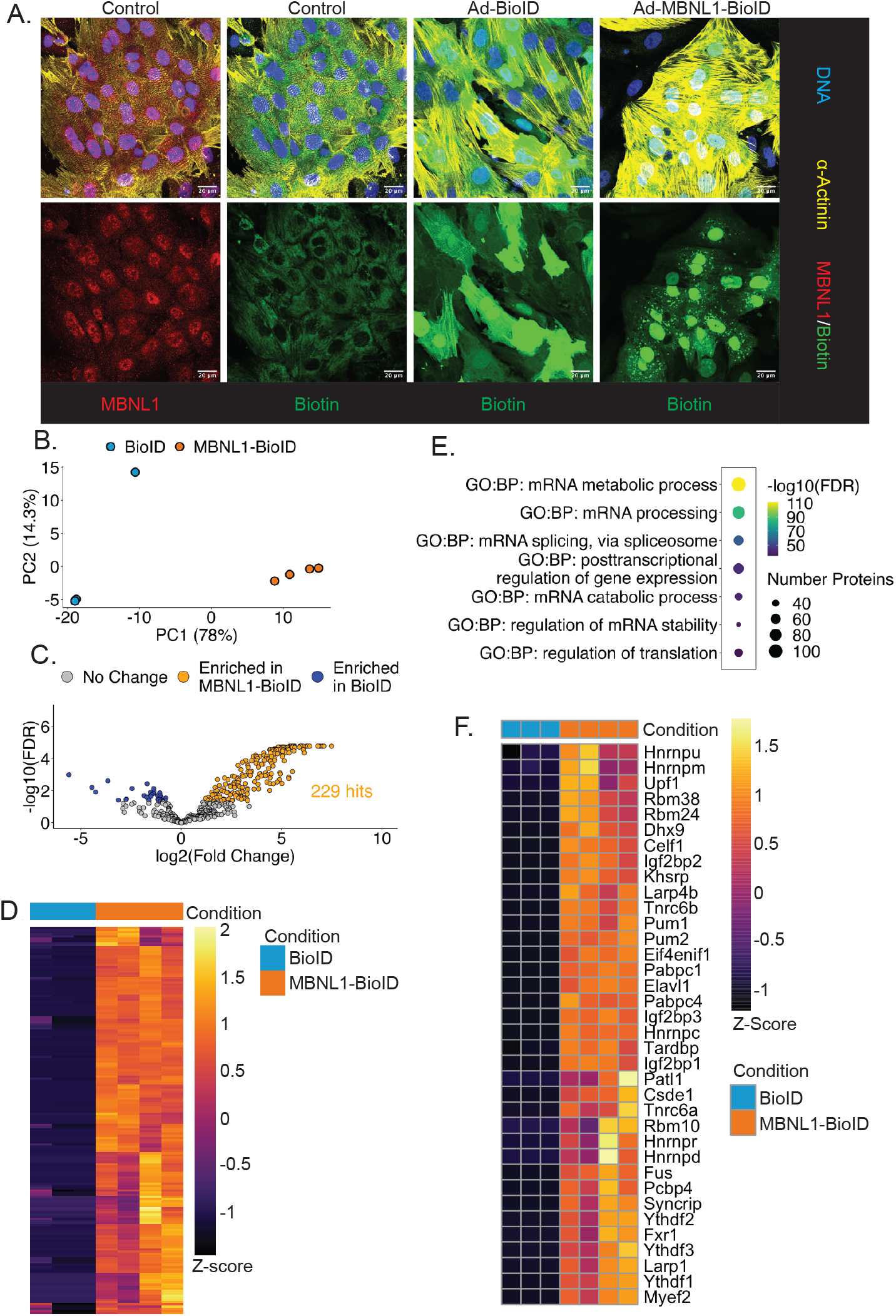
BioID proximity labeling experiment, related to Figure 3. (A) Representative immunocytochemistry images demonstrating the localization of MBNL1 and Biotin in untreated neonatal rat ventricular cardiomyocytes versus those adenovirally transduced with Ad-BioID2 (control) or Ad-MBNL1-BioID2. (B) Principal component analysis of BioID experimental results comparing isolated biotinylated proteins following transduction of unconjugated BioID2 (N=3) or MBNL1-BioID2 (N=4). (C) Volcano plot of differentially enriched proteins. (D) Heatmap of relative enrichment of all proteins significantly enriched in MBNL1-BioID2 (N=4) vs. unconjugated BioID2 control (N=3). (E) Representative Gene Ontology: Biological Processes (GO:BP) terms following gene ontology analysis of proteins enriched in MBNL1-BioID2 vs. unconjugated BioID2 controls. (F) Heatmap of relative enrichment of proteins within the “Regulation of mRNA stability” GO:BP term (N=3/4).

**Supplementary Figure 4.**
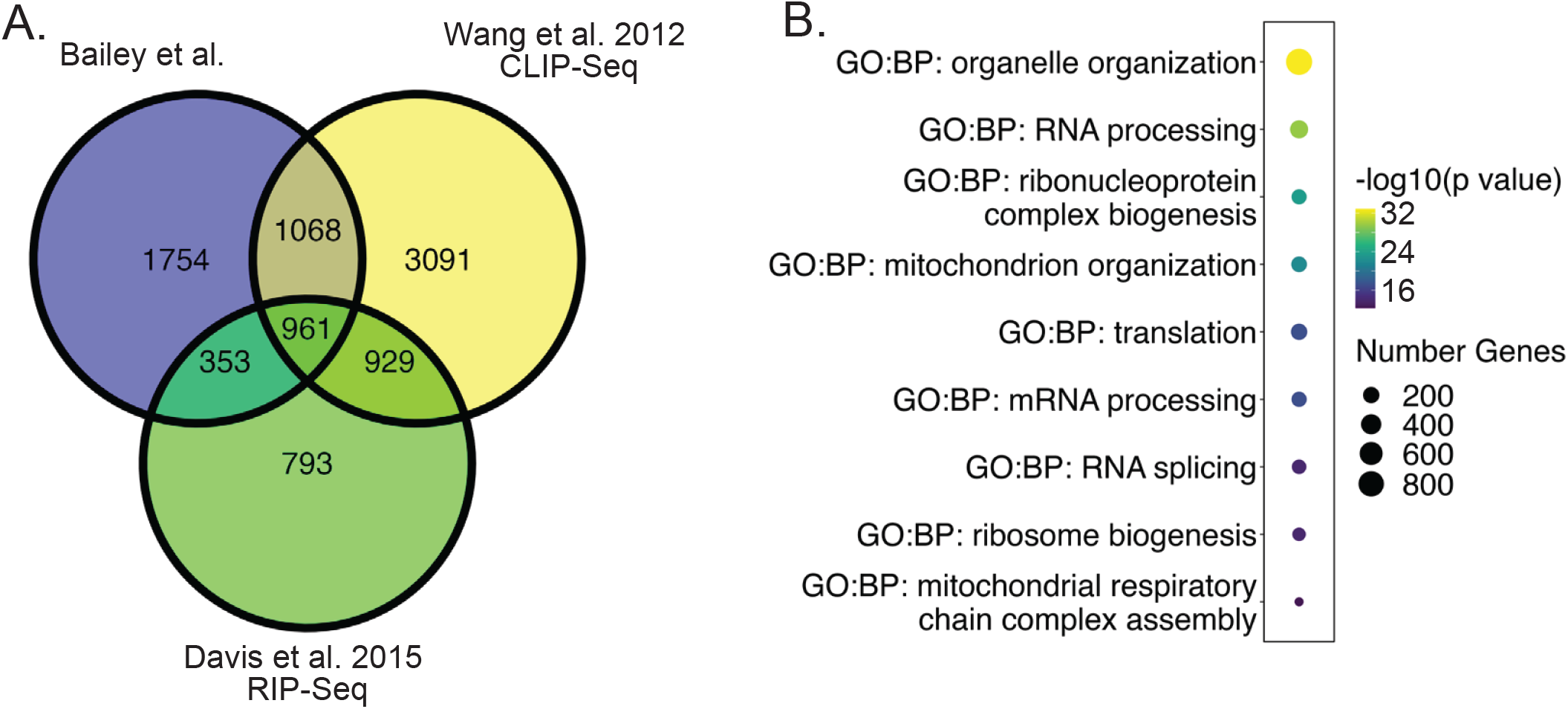
RIPseq analysis, related to Figure 3. (A) Venn diagram comparing MBNL1-bound transcripts identified in this study and two previous studies of MBNL1 target transcripts. (B) Representative GO:BP terms from analysis of input-enriched transcripts from MBNL1 RIPseq.

**Supplementary Figure 5.**
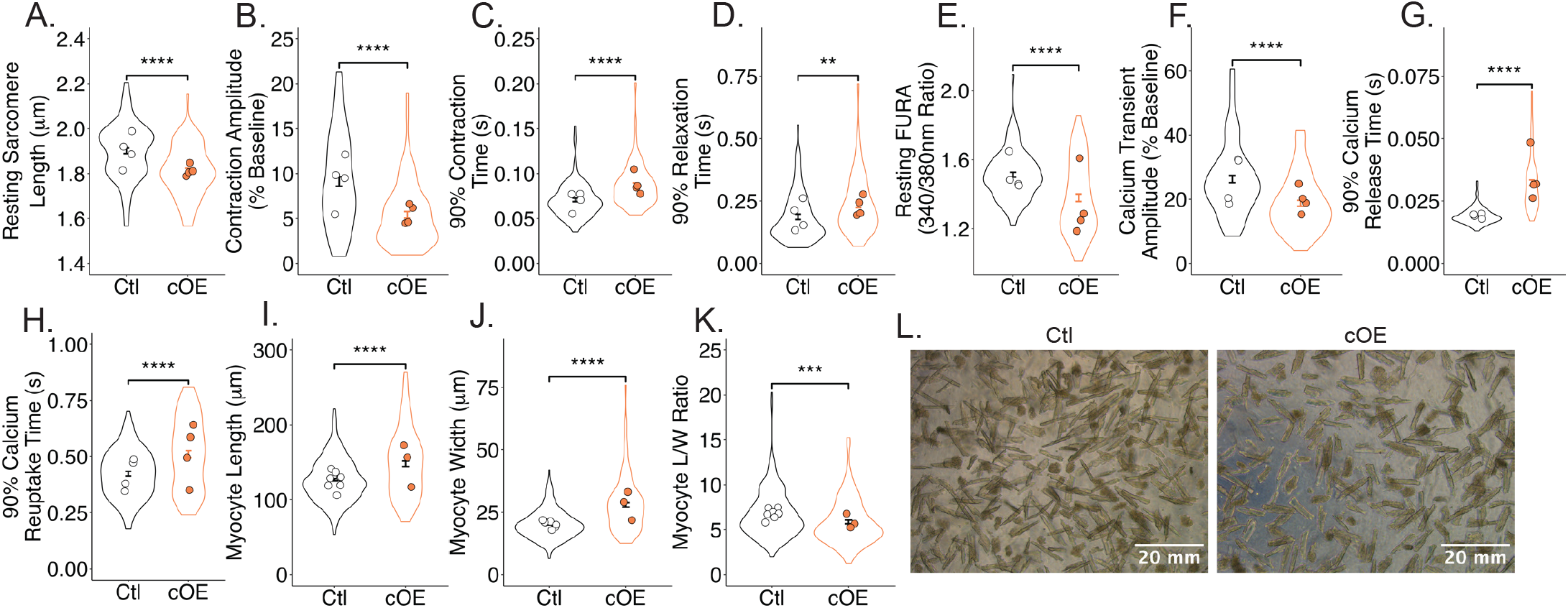
Characterization of cardiomyocytes from control and MBNL1 cOE mice, related to Figure 5. (A-H) Quantification of isolated cardiomyocyte resting sarcomere length (A), contraction amplitude (B), time to 90% contraction (C), time to 90% relaxation (D) resting calcium concentration (FURA ratio) (E), calcium transient amplitude (F), time to 90% calcium release (G), and time to 90% calcium reuptake (H). Dots represent the mean for an individual mouse. Violin plots represent the distribution and error bars show the SEM of individual myocytes. Control n=99 myocytes/N=4 mice and MBNL1 cOE n=85/N=4. **p<0.01,****p<0.0001. (I-K) Quantification of isolated cardiomyocyte length (I), width (J), and length/width ratio (L/W) (K). Dots represent the mean for an individual mouse. Violin plots represent the distribution and error bars show the SEM of individual myocytes. Individual myocytes were compared between groups. Control n=406 myocytes/N=8 mice and MBNL1 cOE n=146/N=3. ***p<0.001, ****p<0.0001. (L) Representative images of control and MBNL1 cOE isolated cardiomyocytes immediately following enzymatic dissociation.

**Supplementary Figure 6.**
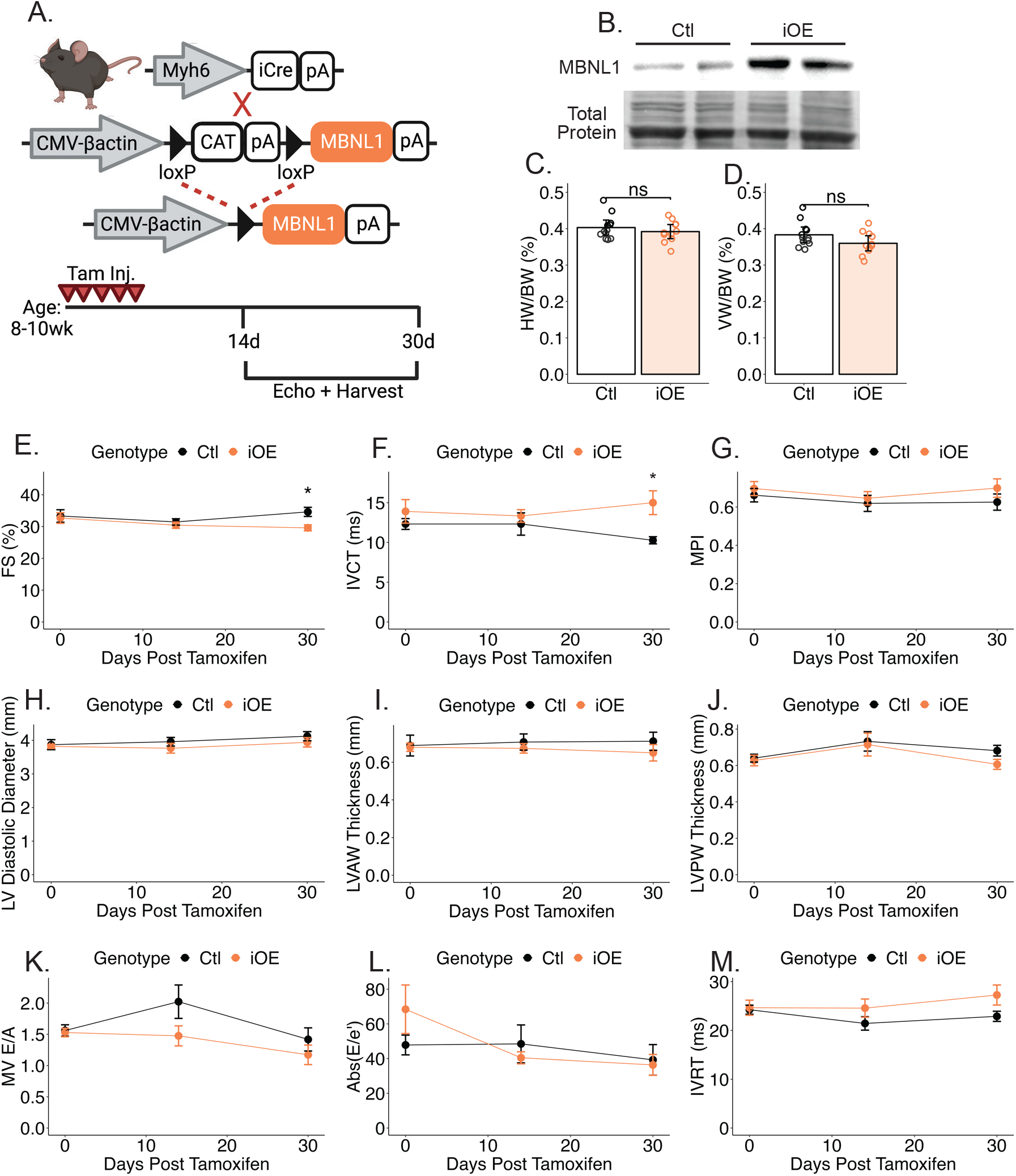
MBNL1 overexpression in adulthood does not recapitulate the phenotype of MBNL1 overexpression in neonatal hearts, related to Figure 5. (A) Breeding schematic to generate MBNL1 iOE mice (top) and experimental schematic (bottom). (B) Western blot for MBNL1 in isolated cardiomyocytes 7-days following tamoxifen initiation. Total protein was used as a loading control (N=2/2). (C-D) Heart weight/body weight (HW/BW) and ventricle weight/body weight (VW/BW) ratios in control and MBNL1 iOE mice 30-days following tamoxifen initiation (N=12/11). Dots represent biological replicates. Bars represent mean ± SEM. ns=not significant. (E-M) Quantification of fractional shortening (FS) (E), isovolumetric contraction time (IVCT) (F), myocardial performance index (MPI) (H), left ventricular (LV) diastolic diameter (H), left ventricular anterior wall (LVAW) diastolic thickness (I), left ventricular posterior wall (LVPW) diastolic thickness (J), mitral valve E/A ratio (MV E/A) (K), absolute value of E/e’ ratio (Abs(E/e’)) (L), and isovolumetric relaxation time (IVRT) (M) by echocardiography (N=8/9). Dots represent mean ± SEM. *p<0.05.

**Supplementary Figure 7.**
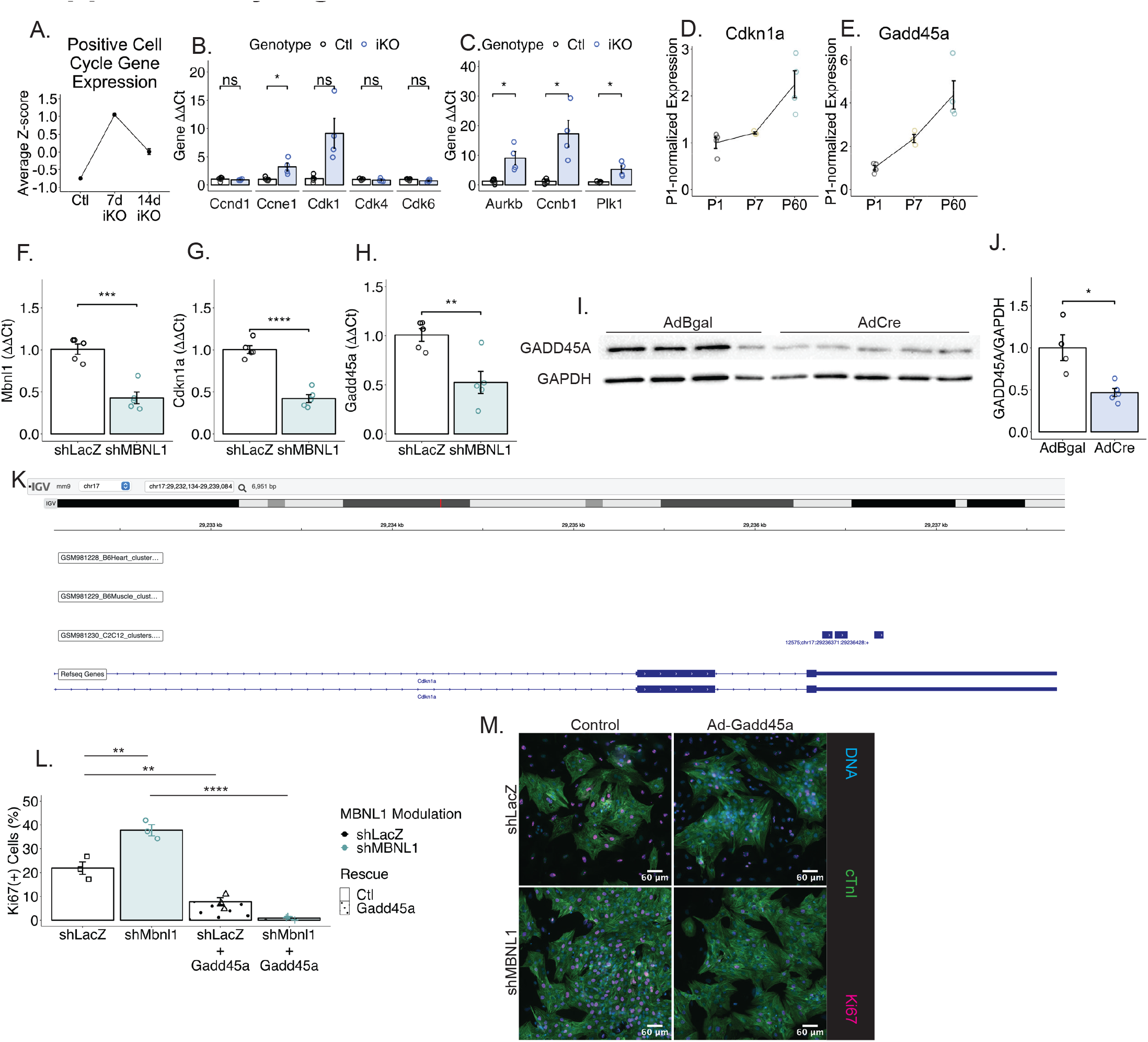
MBNL1 regulates cell cycle inhibitor transcripts, related to Figure 6. (A) Mean normalized expression ± SEM of positive cell cycle regulatory transcripts by RNAseq in control and MBNL1 iKO ventricles. (B-C) Expression of G1/S- (B) and G2/M-relevant (C) cell cycle regulatory genes by qPCR in isolated control and MBNL1 iKO cardiomyocytes (N=5/4). Dots represent biological replicates. Bars represent mean ± SEM. *p<0.05, ns=not significant. (D-E) Mean normalized expression ± SEM of Cdkn1a (D) and Gadd45a (E) by RNAseq over time in wild type ventricles (N=3-4). Dots represent biological replicates. Lines and bars represent mean ± SEM. (F-H) Expression of Mbnl1 (F), Cdkn1a (G), and Gadd45a (H) in shLacZ-versus shMBNL1-treated neonatal rat ventricular cardiomyocytes by qPCR (N=5/5). Dots represent biological replicates. Bars represent mean ± SEM. **p<0.01, ***p<0.001, ****p<0.0001. (I-J) Western blot of GADD45A and GAPDH (I) and quantification of GADD45A (J) in MBNL1^F/F^ mouse neonatal ventricular cardiomyocytes treated with Adβgal or AdCre (N=4/5). Dots represent biological replicates. Bars represent mean ± SEM. *p<0.05. (K) Integrated Genomics Viewer (IGV) screenshot of location of MBNL1 CLIP clusters in the 3’UTR of the Cdkn1a transcript. (L-M) Quantification (L) and representative images (M) of cardiomyocyte Ki67-positivity in adenovirus-treated neonatal rat ventricular cardiomyocytes (N=3). Dots represent biological replicates. Bars represent mean ± SEM. **p<0.01, ****p<0.0001.

**Table S1.**
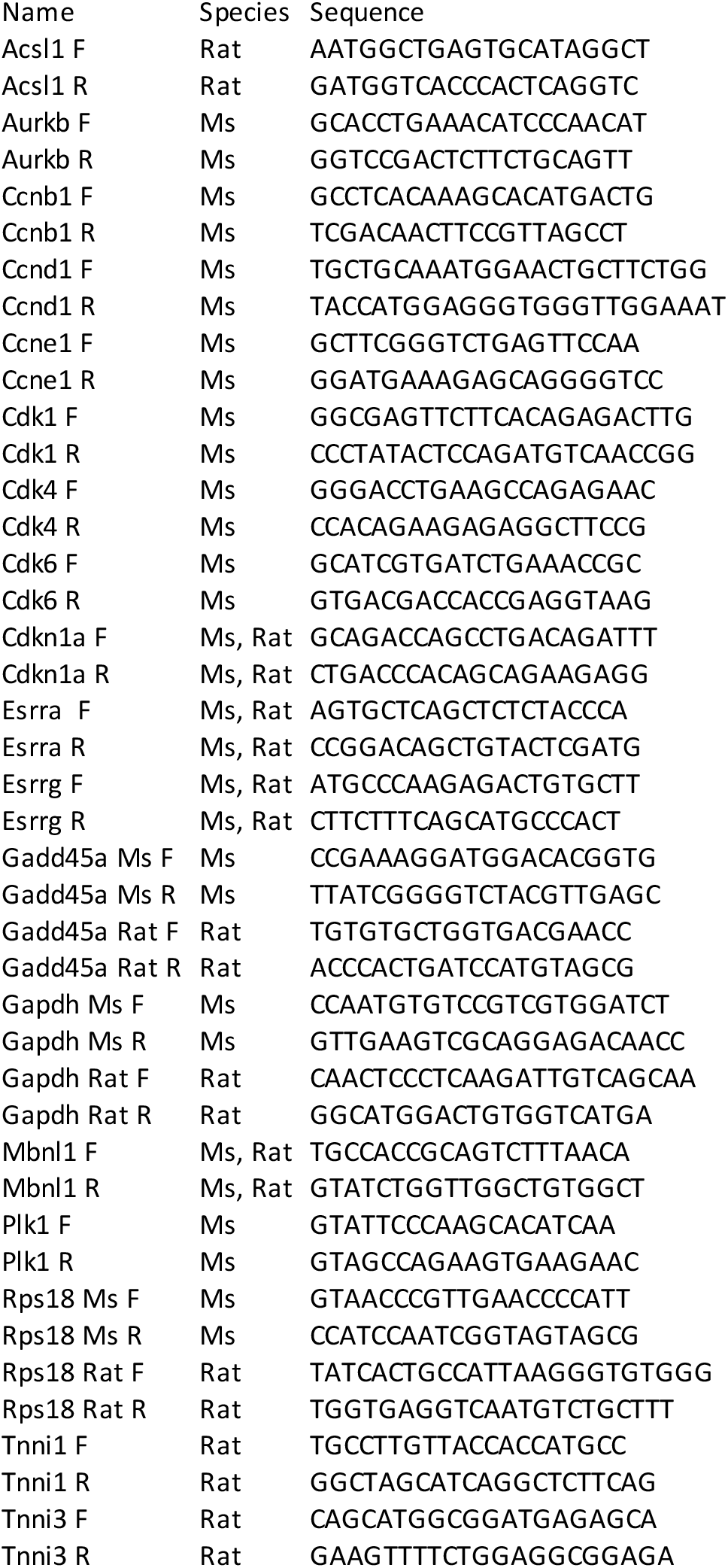

**Table S2.**
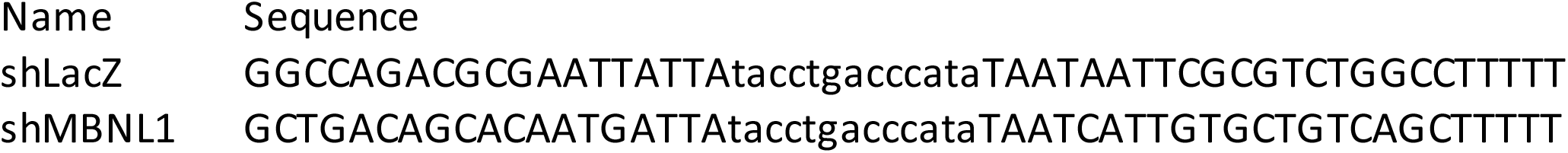

## Notes

### Competing Interest Statement

The authors have declared no competing interest.

